# PI3P dependent regulation of cell size and autophagy by phosphatidylinositol 5-phosphate 4-kinase

**DOI:** 10.1101/2022.06.18.496676

**Authors:** Avishek Ghosh, Aishwarya Venugopal, Dhananjay Shinde, Sanjeev Sharma, Meera Krishnan, Swarna Mathre, Harini Krishnan, Padinjat Raghu

**Affiliations:** National Centre for Biological Sciences, TIFR-GKVK Campus, Bellary Road, Bangalore 560065, India

## Abstract

Phosphatidylinositol 3-phosphate (PI3P) and Phosphatidylinositol 5-phosphate (PI5P) are low abundant phosphoinositides crucial for key cellular events such as endosomal trafficking and autophagy. Phosphatidylinositol 5-phosphate 4-kinase (PIP4K) is an enzyme that regulates PI5P *in vivo* but can act on both PI5P and PI3P, *in vitro*. In this study, we report a novel role for PIP4K in regulating PI3P levels in *Drosophila* tissues. Loss-of-function mutants of the only PIP4K gene in *Drosophila* (*dPIP4K^29^*) show reduced cell size in larval salivary glands. We find that PI3P levels are elevated in *dPIP4K^29^* tissues and that reverting PI3P levels back towards wild type, without changes in PI5P levels, can also rescue the reduced cell size phenotype. *dPIP4K^29^* mutants also show an upregulation in autophagy and the reduced cell size can be reverted by decreasing Atg8a, that is required for autophagosome maturation. Lastly, increasing PI3P levels in wild type salivary glands can phenocopy the reduction in cell size and associated upregulation of autophagy seen in *dPIP4K^29^*. Thus, our study reports for the first time, a role for a PIP4K-regulated PI3P pool in the control of autophagy and cell size regulation that may explain the reported role of PIP4K in regulating neurodegeneration and tumour growth.

## Introduction

The organization of membranes in eukaryotic cells is regulated by signalling mechanisms that couple ongoing stimuli to sub-cellular transport mechanisms. Several signalling molecules contribute to this process including proteins such as SNAREs and Rabs along with lipids. Phosphoinositides are a class of signalling lipids found in all eukaryotes; they are glycerophospholipids whose inositol headgroup can be phosphorylated on the 3^rd^, 4^th^ or 5^th^ positions to generate molecules with signalling functions (Balla, 2013). In cells, phosphoinositides are generated by the action of lipid kinases that are able to add phosphate groups to specific positions on the inositol head group of specific substrates (Sasaki et al., 2009); thus the activity of these lipid kinases and phosphatases is important to generate lipid signals on organelle membranes. Phosphatidylinositol 5 phosphate 4-kinase (PIP4K) are one such class of lipid kinases that convert phosphatidylinositol 5 phosphate (PI5P) into phosphatidylinositol 4,5 bisphosphate [PI(4,5)P_2_] (Clarke and Irvine, 2013; Rameh et al., 1997). Genetic analysis of *PIP4K* in various organisms have demonstrated their importance in development and growth control (Gupta et al., 2013), cell division (Emerling et al., 2013) immune cell function (Shim et al., 2016), metabolism (Lamia et al., 2004) and neurological disorders (Al-Ramahi et al., 2017). At a cellular level, PIP4K have been implicated in the control of plasma membrane receptor signalling (Sharma et al., 2019), vesicular transport (Kamalesh et al., 2017), autophagy (Lundquist et al., 2018; Vicinanza et al., 2015) and nuclear functions such as transcriptional control (Fiume et al., 2019). PI(4,5)P_2_, the product of PIP4K activity has many important functions in regulating cell physiology and signalling (Kolay et al., 2016) and PI5P, the well-defined substrate of PIP4K has also been implicated in regulating some sub-cellular processes (Hasegawa et al., 2017). However, despite their importance in regulating several cellular processes and physiology, the biochemical reason for the requirement of PIP4K in regulating these processes remain unclear.

When studied using biochemical activity assays *in vitro*, PIP4K shows very high activity on PI5P to generate PI(4,5)P_2_ (Ghosh et al., 2019; Rameh et al., 1997; Zhang et al., 1997). Coupled with this, analysis of lipid levels following genetic depletion of PIP4K in various models have failed to note appreciable reductions in PI(4,5)P_2_ levels [reviewed in (Kolay et al., 2016)]. Rather, such studies have reported an increase in the levels of the substrate, PI5P (Gupta et al., 2013; Jones et al., 2006; Stijf-Bultsma et al., 2015) suggesting that the relevant biochemical function of the enzyme is to regulate PI5P levels. Previous studies have noted that PIP4K depletion in *Drosophila* photoreceptors (Kamalesh et al., 2017) leads to altered endocytic function and a role for PI5P in regulating endocytosis has been proposed (Boal et al., 2015; Ramel et al., 2011). In mammalian cells, PI5P has been proposed as a mediator of autophagy regulation by PIP4K (Al-Ramahi et al., 2017; Lundquist et al., 2018; Vicinanza et al., 2015). PIP4K can also utilise PI3P as a substrate *in vitro* (Ghosh et al., 2019; Gupta et al., 2013; Zhang et al., 1997), albeit with low efficiency; however, the significance of this activity *in vivo* and the role of PIP4K, if any in regulating PI3P levels *in vivo* is not known. PI3P is well known as a regulator of autophagy (Schink et al., 2016; Wallroth and Haucke, 2018), a process that is reported to be altered on modulating PIP4K function but the significance, if any, of PIP4K regulated pools of PI3P in these processes remains unknown. PI3P formed at the phagophore membrane by Vps34, a class III PI3-kinase is important for autophagy initiation by recruiting proteins like DFCP1, WIPI (Axe et al., 2008; Polson et al., 2010). In the next step, the ATG16L1 complex, which includes the proteins ATG16L1, ATG5 and ATG12, is recruited to the pre-autophagosomal membranes (Dudley et al., 2019)and Myotubularins, 3-phosphatase enzymes that have been reported to regulate autophagy by regulating PI3P levels at autophagy initiation membranes (Taguchi-Atarashi et al., 2010; Vergne et al., 2009; Zou et al., 2012). This raises the possibility that the reported regulation of autophagy by PIP4K may arise from its ability to regulate PI3P levels at the autophagic membrane?

The *Drosophila* genome contains a single gene encoding PIP4K (*dPIP4K*). A loss-of-function allele of dPIP4K (*dPIP4K^29^)* results in altered growth and development, accumulation of the known substrate PI5P and no reduction in PI(4,5)P_2_ levels (Gupta et al., 2013). In *dPIP4K^29^*, the size of larval salivary gland cells is reduced and genetic reconstitution studies have demonstrated that the kinase activity of dPIP4K, is required to support normal cell size (Mathre et al., 2019). Previous work has shown that TORC1 signalling, a known regulator of cell size (Lloyd, 2013) and autophagy (Nascimbeni et al., 2017), is reduced in *dPIP4K^29^* (Gupta et al., 2013). Thus, while it is clear that the kinase activity of dPIP4K is required for normal salivary gland cell size, the biochemical basis for this requirement of enzyme activity is not known.

In this study, we show that in addition to the previously reported elevation of PI5P levels, PI3P levels are also elevated in *dPIP4K^29^* and this elevation in PI3P is dependent on the kinase activity of dPIP4K. The reduced salivary gland cell size in *dPIP4K^29^* could be rescued by the expression of a PI3P specific 3-phosphatase, *Mtm* and this rescue was associated with a reversal of the elevated PI3P but not PI5P levels. Interestingly, we observed that in larval salivary glands of *dPIP4K^29^,* the elevation in PI3P levels was associated with an upregulation in autophagy and the phenotype of reduced cell size in *dPIP4K^29^* could be reversed by down-regulating Atg8a, which functions downstream to the formation of PI3P in the autophagy pathway. Elevation of PI3P levels in wild type salivary glands by depletion of *Mtm* resulted in both a reduction in cell size and the enhanced autophagy in salivary glands. Therefore, this study underscores a novel *in vivo* regulation of PI3P levels by PIP4K in a multicellular organism leading to the control of cell size.

## Results

### *dPIP4K* does not regulate cell size through levels of its product PI(4,5)P_2_

The kinase activity of dPIP4K is required for its ability to support salivary gland cell size (Figure 1A depicts the conversion of PI5P to PI(4,5)P_2_ by dPIP4K) (Mathre et al., 2019). Thus its ability to regulate cell size may depend either on the elevated levels of its preferred substrate PI5P, or a shortfall in the pool of the product PI(4,5)P_2_ generated. Previous studies have identified a point mutation (A381E) in PIP4Kβ that can switch its substrate specificity from PI5P to PI4P while still generating the same product PI(4,5)P_2_ (Kunz et al., 2002). The corresponding point mutant version of hPIP4Kα has been used to distinguish between phenotypes dependent on the PI(4,5)P_2_ generated by PIP4K as opposed to PI5P metabolised by it (Bulley et al., 2016). We generated a switch mutant version of human PIP4Kβ, hPIP4Kβ^[A381E]^ that cannot utilise PI5P as a substrate but can produce PI(4,5)P_2_ using PI4P as a substrate (Kunz et al., 2002). Expression of hPIP4Kβ^[A381E]^ in the salivary glands of *dPIP4K^29^* (AB1> hPIP4Kβ^[A381E]^; *dPIP4K^29^*) (Figure S1A) did not rescue the reduced cell size whereas reconstitution with the wild type enzyme was able to do so as previously reported (Figure S1Bi, quantified in Figure S1Bii, western blot in Figure S1A) (Mathre et al., 2019). This observation suggests that the ability of dPIP4K to regulate cell size does not depend on the pool of PI(4,5)P_2_ that it generates and also that regulation of the levels of its substrate is likely to be the relevant biochemical basis through which the enzyme supports cell size in salivary glands.

**Figure 1:**
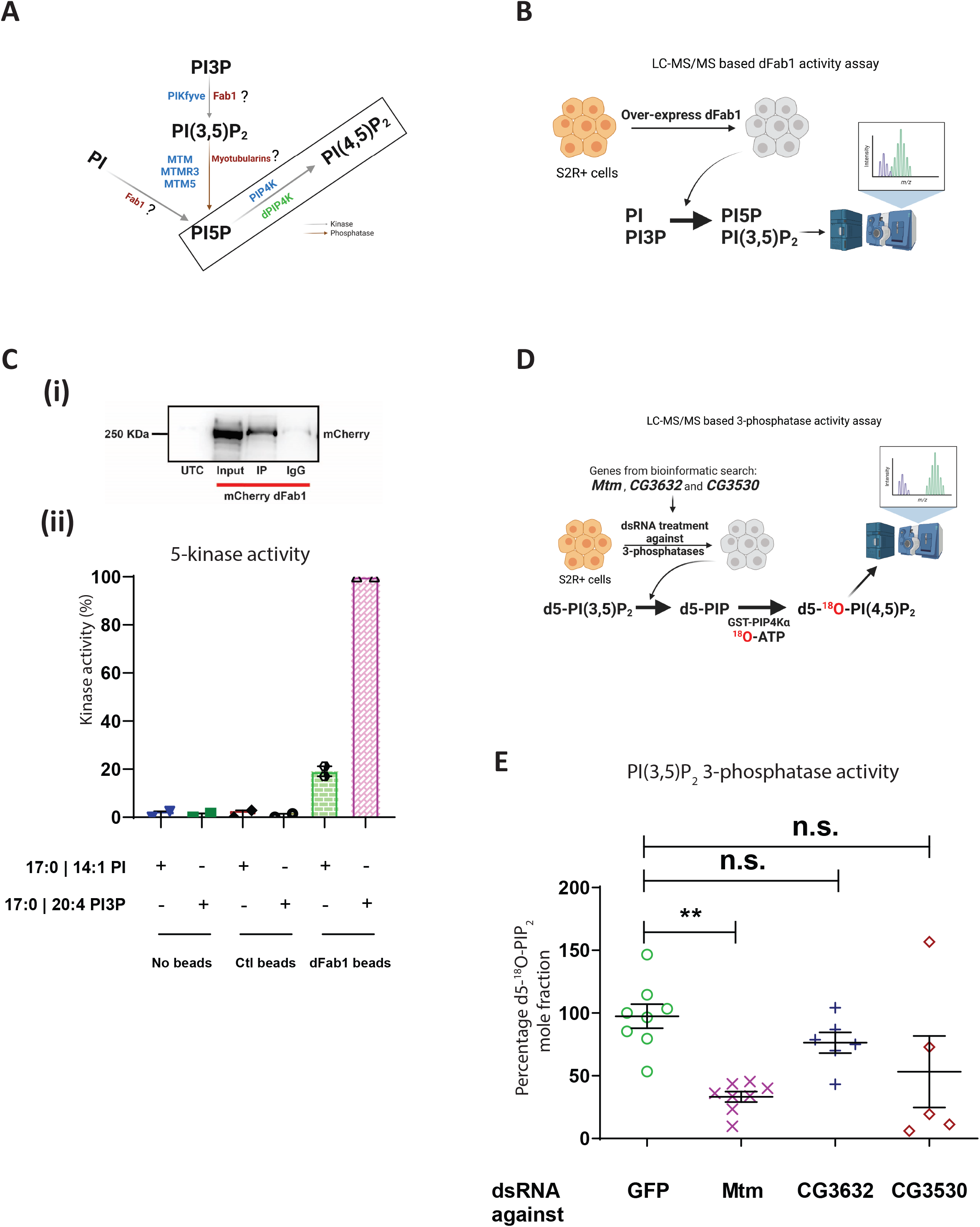
Screening for a biochemical route to modulate PI5P levels in *Drosophila* as altering local PI(4,5)P_2_ couldn’t change cell size of *dPIP4K^29^*. (A) Schematic illustrating the putative enzymatic routes by which PI5P can be synthesised in *Drosophila*. The activities of enzymes labelled in blue are known in mammalian cells, the activity of enzymes labelled in red followed by “?” are still unknown in *Drosophila*, the activity of enzymes labelled in green are known in *Drosophila*. The activity of PI5P to PI(4,5)P_2_ is boxed and is linked to cell size regulation in *Drosophila* (Mathre et al., 2019). PI: Phosphatidylinositol, PI3P: Phosphatidylinositol 3-phosphate, PI(3,5)P_2_: Phosphatidylinositol 3,5 bisphosphate, PI5P: Phosphatidylinositol 5-phosphate, PI(4,5)P_2_: Phosphatidylinositol 4,5 bisphosphate. (B) Schematic illustrating the LC-MS/MS based *in vitro* 5-kinase activity assay using S2R+ cells over-expressing dFab1 enzyme to convert synthetic PI or PI3P to PI5P and PI(3,5)P_2_, respectively. (C) (i) Immunoprecipitated protein levels were analysed by Western blotting with an anti-mCherry antibody. Control (IgG) was prepared without anti-mCherry. Input lane shows the correct size of dFab1 protein ∼230 kDa. UTC: Untransfected control. (ii) *In vitro* kinase assay on synthetic PI and PI3P. Graph representing the Kinase activity (%) as the normalised response ratio of “PI 5-kinase activity on PI” to “PI3P 5-kinase activity on PI3P” upon enzymatic activity of immunoprecipitated mCherry::dFab1 on the respective substrates. Response ratio of PI 5-kinase activity on PI is obtained from area under the curve (AUC) of 17:0 14:1 PI5P (Product)/17:0 14:1 PI (Substrate), Response ratio of PI3P 5-kinase activity on PI3P is obtained from area under the curve (AUC) of 17:0 20:4 PI(3,5)P_2_ (Product)/17:0 20:4 PI3P (Substrate) and is represented as mean ± S.E.M. on addition of either negative control (no beads), Control (mCherry beads) or dFab1 (mCherry::dFab1 beads). Number of immunoprecipitated samples = 2. (D) Schematic illustrating the LC-MS/MS based *in vitro* PI(3,5)P_2_ 3-phosphatase activity assay using dsRNA treated S2R+ cells as enzyme source to convert synthetic PI(3,5)P_2_ [d5-PI(3,5)P_2_ to d5-^18^O-PIP_2_] using a two-step reaction scheme. (E) *In vitro* phosphatase assay on synthetic PI(3,5)P_2_. Graph representing the 3-Phosphatase activity (%) as the percent formation of d5-^18^O-PIP_2_ formed from starting d5-PI(3,5)P_2_ as mean ± S.E.M. on addition of either control (GFP ds RNA) or Mtm, CG3632, CG3530 ds RNA treated S2R+ cell lysates. One way ANOVA with a post hoc Tukey’s test shows p value = 0.003 between GFP and Mtm ds RNA treated lysates, shows p value = 0.63 between GFP and CG3632 ds RNA treated lysate and shows p value = 0.11 between GFP and CG3530 ds RNA treated lysates.

### *Mtm* could be a candidate gene to modulate PI5P levels in *Drosophila*

Since PI5P is the preferred substrate of dPIP4K (Gupta et al., 2013), we sought to modulate PI5P levels to assess the impact on cell size regulation. However, other biochemical players involved in PI5P regulation in *Drosophila* are unknown so far. In mammals, PI5P levels are regulated by PIKFYVE, the type III PIP 5-kinase that converts PI3P to PI(3,5)P_2_ and PI to PI5P (Hasegawa et al., 2017; Shisheva, 2013). *Drosophila* has a single PIKFYVE homologue (*CG6355*, here named *dFab1*) (Rusten et al., 2006); however, its biochemical activity has not be tested (Figure 1A). We expressed mCherry tagged dFab1 in S2R+ cells, immuno-precipitated it (Figure 1Ci) and analysed its ability to phosphorylate PI3P and PI, using a LC-MS/MS based *in vitro* kinase activity assay for dFab1 (Figure 1B shows a schematic for the assay). We found that the relative activity of dFab1 on synthetic PI3P was approximately 4 times greater than the activity on synthetic PI (Figure 1Cii). Since dFab1 preferentially synthesizes PI(3,5)P_2_ from PI3P, subsequent PI5P generation would require the activity of a 3-phosphatase that can dephosphorylate PI(3,5)P_2_. In mammals, *in vitro* studies have revealed that lipid phosphatases of the myotubularin family have specific activity toward PI3P and PI(3,5)P_2_ (Figure 1A) (Laporte et al., 1996; Schaletzky et al., 2003; Walker et al., 2001). In most higher order organisms, there are multiple myotubularin isoforms (Robinson and Dixon, 2006). It has been suggested that *Drosophila* has six isoforms (Oppelt et al., 2013), but bioinformatic analysis using multiple sequence alignment revealed that the conserved CX_5_R catalytic motif is present in only 3 genes – *Mtm*, *CG3632* and *CG3530* (Figure S1C). To identify the myotubularin that might generate PI5P from PI(3,5)P_2_, we designed a two-step *in vitro* LC-MS/MS based PI(3,5)P_2_ 3-phosphatase assay using *Drosophila* S2R+ cell lysates as a source of enzyme (Figure 1D, details of the assay is mentioned in methods). Briefly, deuterium labelled PI(3,5)P_2_ [d5-PI(3,5)P_2_] is incubated with cell lysate and the PI5P formed through the action of a 3-phosphatase is converted, using ^18^O-ATP to PI(4,5)P_2_ of an unique mass owing to the incorporated ^18^O, and subsequently detected on a mass spectrometer (Figure 1D). We used a linked PI5P-4-kinase assay to distinguish a 3-phosphatase activity generating PI5P from a 5-phosphatase activity generating PI3P, from the cell lysates in the first step of the assay. Each of the 3-phosphatases (*Mtm*, *CG3632* and *CG3530*) were depleted using dsRNA treatment (Worby et al., 2001) and the 3’ phosphatase activity of the lysates were measured. We noted more than 50% knockdown using dsRNA against *Mtm*, *CG3632* and *CG3530* in S2R+ cells (Figure S1Di-iii). We observed that the d5-^18^O-PIP_2_ mole fraction [the measure of PI(3,5)P_2_ 3-phosphatase activity] for *Mtm* downregulated lysates was significantly lower as compared to control GFP dsRNA treated lysates (Figure 1E). However, we did not observe a significant difference in activity of lysates downregulated for *CG3632* or *CG3530*. Therefore, Mtm is a 3-phosphatase that could regulate PI5P synthesis in *Drosophila*.

### *Drosophila* Mtm reverses the cell size defect of *dPIP4K^29^* independent of PI5P levels

Based on the results of our *in vitro* results, we over-expressed *Mtm* in *dPIP4K^29^* salivary glands to elevate PI5P levels. If the reduced cell size in *dPIP4K^29^* was linked to elevated PI5P levels, *Mtm* over-expression in *dPIP4K^29^* is expected to lead to a further reduction in cell size. Surprisingly, we observed that over-expression of *Mtm* in the salivary glands of *dPIP4K^29^* (*AB1> MtmGFP*; *dPIP4K^29^*) resulted in a reversal of cell size as compared to *dPIP4K^29^* glands (Figure 2Ai, quantified in Figure 2Aii); over-expression of the enzyme in wild type salivary glands did not affect cell size (Figure S2A).

**Figure 2:**
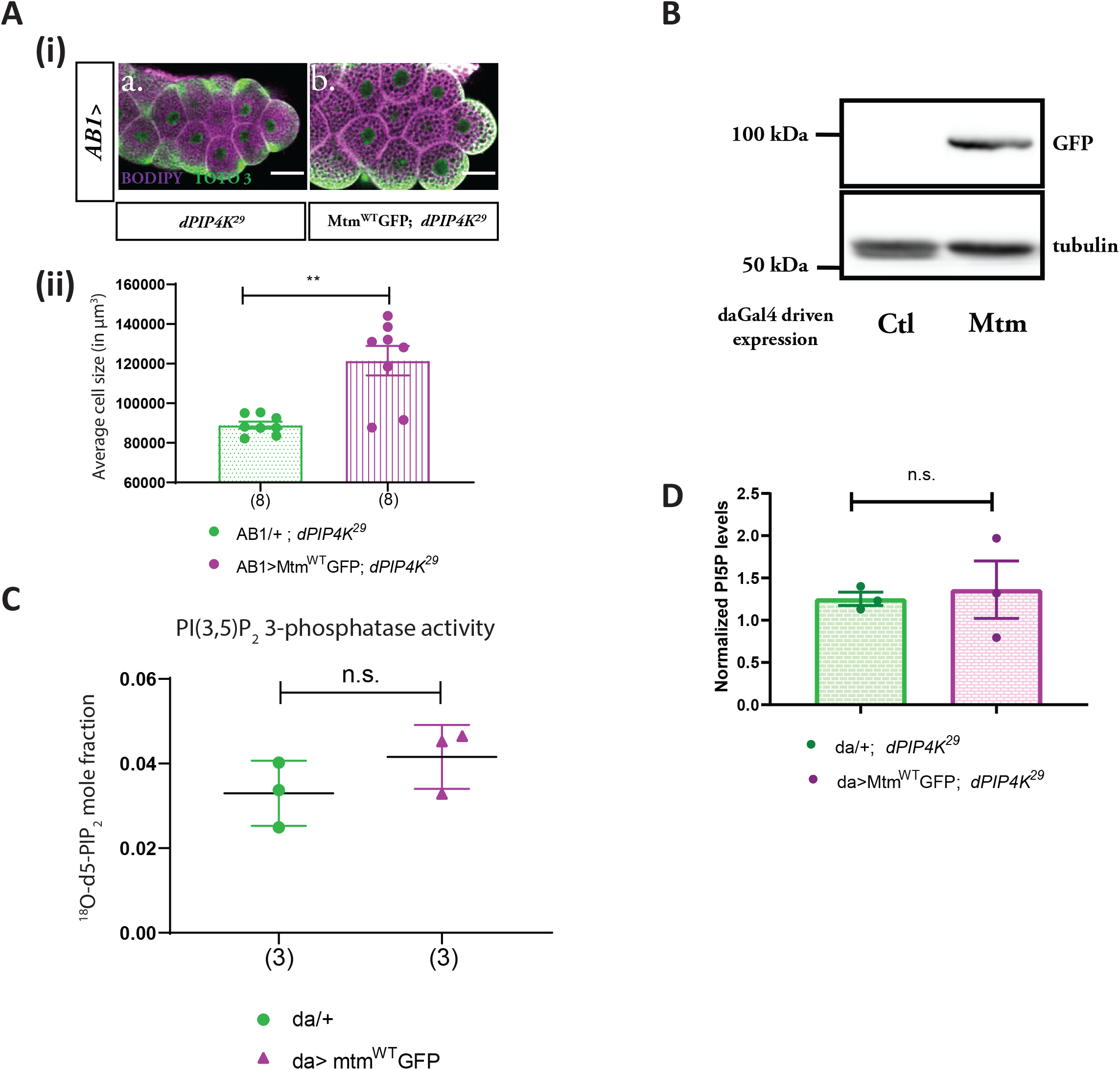
*Drosophila* Mtm rescues the cell size defect of *dPIP4K^29^* independent of PI5P levels. (A) (i) Representative confocal images of salivary glands from the genotypes a. *AB1/+; dPIP4K^29^*, b. *AB1>*Mtm^WT^GFP *; dPIP4K^29^*. Cell body is marked majenta by BODIPY conjugated lipid dye, nucleus is marked by TOTO-3 shown in green. Scale bar indicated at 50 μm. (ii) Graph representing average cell size measurement (in percentage) as mean ± S.E.M. of salivary glands from wandering third instar larvae of *AB1/+; dPIP4K^29^* (n = 8), *AB1>*Mtm^WT^GFP*; dPIP4K^29^* (n = 8). Sample size is represented on individual bars. Student’s unpaired t-test with Welch correction showed p value = 0.003 between *AB1/+; dPIP4K^29^* and *AB1>*Mtm^WT^GFP *; dPIP4K^29^*. (B) Protein levels between *daGal4*/+ (Ctl) and *da*> Mtm^WT^GFP from third instar wandering larvae seen on a Western blot probed by GFP antibody. Mtm^WT^GFP migrates ∼100 kDa. Tubulin was used as the loading control. (C) *In vitro* phosphatase assay on synthetic PI(3,5)P_2_. Graph representing the formation of ^18^O-PIP_2_ formed from starting PI(3,5)P_2_ as substrate represented as mean ± S.E.M. on addition of either control (da/+) or da>Mtm_GFP lysates. Lysate samples n = 3, where each sample was made from five third instar wandering larvae. Student’s unpaired t-test with Welch correction showed p value = 0.23. (D) Graph representing Normalised PI5P levels which is total ^18^O-PIP_2_/peak area of 17:0 20:4 PI(4,5)P_2_ (internal standard) normalised to organic phosphate value as mean ± S.E.M. of da/+; *dPIP4K^29^* (green) or da> Mtm^WT^GFP, *dPIP4K^29^* (majenta). Biological samples n = 3, where each sample was made from five third instar wandering larvae. Unpaired t test with Welch’s correction showed p value = 0.7830 between da/+; *dPIP4K^29^* and da> Mtm^WT^GFP, *dPIP4K^29^*.

Mtm is a 3-phosphatase that can act on PI3P to produce PI and PI(3,5)P_2_ to produce PI5P. Previously, its activity on PI(3,5)P_2_ has been demonstrated using purified protein in an *in vitro* phosphate-release assay (Velichkova et al., 2010). To understand the biochemical basis of the ability of over-expressed Mtm to reverse cell size in *dPIP4K^29^*, we tested the biochemical activity of Mtm from *Drosophila* larval extracts using our two-step 3-phosphatase activity assay. Figure 2B shows expression of C-terminus GFP tagged Mtm from larval lysates at molecular weights as predicted *in silico*. We found that overexpression of Mtm did not result in a statistically significant increase in 3-phosphatase activity compared to controls (Figure 2C). To confirm this result was not a result of C-terminal tagging leading to Mtm inactivation, we cloned an N-terminus mCherry tagged Mtm and performed the 3-phosphatase assay using S2R+ cell lysates expressing mCherry_Mtm (Figure S2Bi). It was observed that a N-terminally tagged Mtm was also not active on PI(3,5)P_2_ as compared to controls, much like its C-terminal GFP tagged counterpart (Figure S2Bii). These findings suggest that the generation of PI5P from PI(3,5)P_2_ by Mtm in *Drosophila* larvae is likely to be minimal. We also measured the levels of PI5P from larval lipid extracts using a recently standardised LC-MS/MS based PI5P mass assay (Ghosh et al., 2019), comparing larvae expressing Mtm in *dPIP4K^29^* mutant background with *dPIP4K^29^*. We observed that the overexpression of Mtm did not alter the levels of PI5P in *dPIP4K^29^* (Figure 2D). Therefore, together we conclude that (a) Mtm cannot synthesise PI5P from PI(3,5)P_2_ *in vivo* in *Drosophila* and (b) Mtm expression rescued the cell size of *dPIP4K^29^* without changing the elevated PI5P levels. Therefore, we investigated PI5P independent mechanism that control cell size.

### Mtm reduces PI3P levels when over-expressed in *dPIP4K^29^*

Mtm has also been shown to dephosphorylate PI3P to generate PI *in vitro* (Velichkova et al., 2010). We tested the activity of lysates expressing Mtm on synthetic PI3P using a LC-MS/MS based assay and found that lysates with Mtm over-expression showed significantly higher PI3P 3-phosphatase activity compared to control lysates (Figure 3A), raising the possibility that Mtm might be rescuing cell size in *dPIP4K^29^* through its PI3P 3-phosphatase activity.

**Figure 3:**
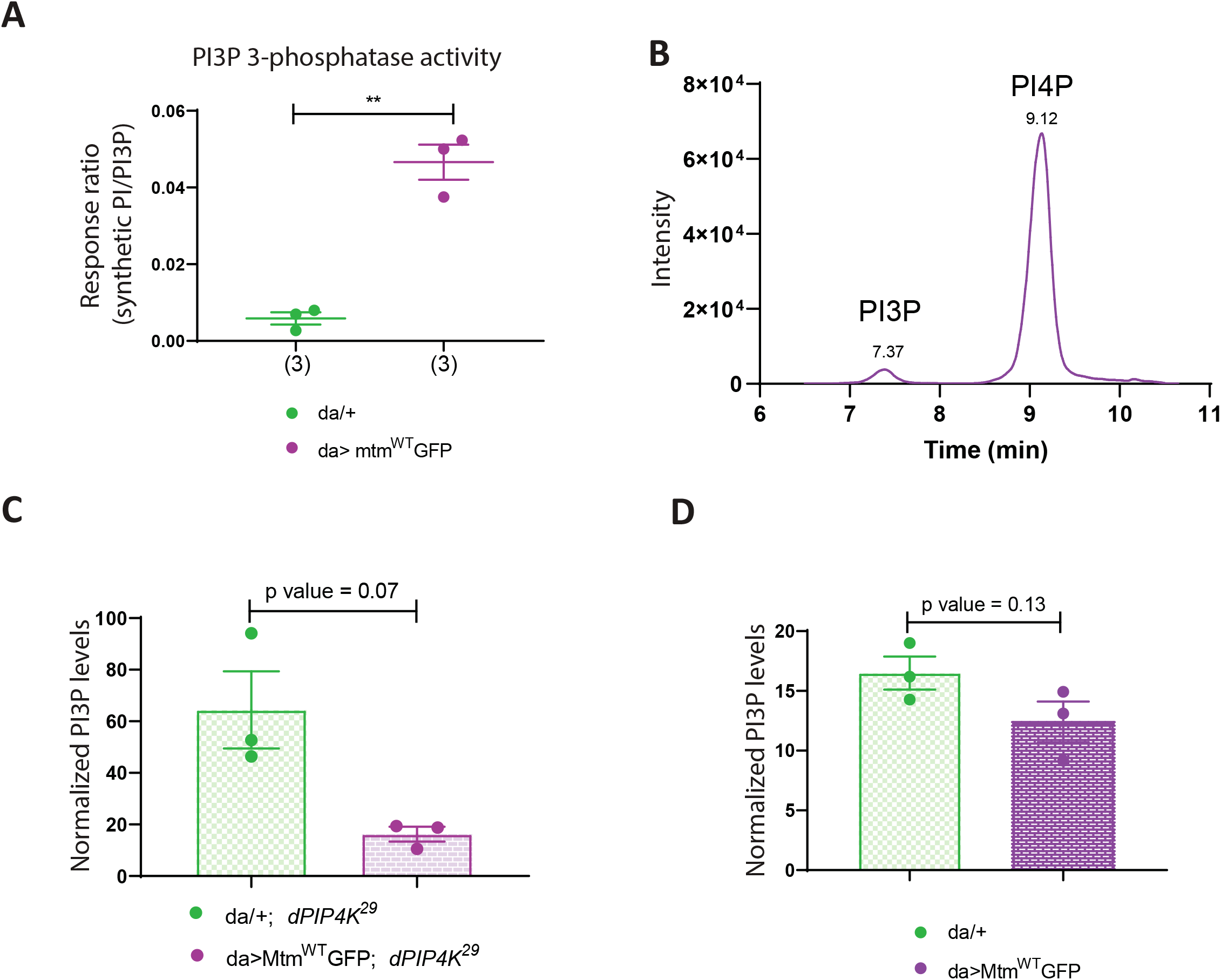
Mtm reduces PI3P levels when over-expressed in *dPIP4K^29^*. (A) *In vitro* phosphatase assay on synthetic PI3P. Graph representing the response ratio of 17:0 20:4 PI (Product)/17:0 20:4 PI3P (Substrate) formed as mean ± S.E.M. on addition of either control (da/+) or da>Mtm^WT^GFP lysates. Lysate samples = 3, where each sample was made from five third instar wandering larvae. Student’s unpaired t-test with Welch correction showed p value = 0.007. (B) Extracted ion chromatogram (XIC) of deacylated PI3P or GroPI3P (Glycerophosphoinositol 3-phosphate) peak at Rt = 7.37 min, separated from deacylated PI4P or GroPI4P (Glycerophosphoinositol 4-phosphate) peak at Rt = 9.12 min obtained from injecting wild type larval lipid extract (details of sample preparation is discussed in methods). (C) Graph representing Normalised PI3P levels which is the peak area of GroPI3P/ peak area of GroPI4P normalised to organic phosphate value of total lipid extracts as mean ± S.E.M. of da/+ ; *dPIP4K^29^* (green) and da> Mtm^WT^GFP, *dPIP4K^29^* (majenta). Biological samples = 3, where each sample was made from three third instar wandering larvae. Student’s unpaired t-test with Welch correction showed p value = 0.07. (D) Graph representing Normalised PI3P levels which is the peak area of GroPI3P/ peak area of GroPI4P normalised to organic phosphate value of total lipid extracts as mean ± S.E.M. of da/+ (green) and da> Mtm^WT^GFP, (majenta). Biological samples = 3, where each sample was made from three third instar wandering larvae. Student’s unpaired t-test with Welch correction showed p value = 0.13.

Mtm activity can in principle change the levels of PI and PI3P; however since PI3P levels in cells are substantially lower (<10%) of PI (Stephens et al., 1993), we analysed PI3P levels in relation to the ability of Mtm overexpression to rescue the reduced cell size in *dPIP4K^29^* salivary glands. Currently used methods to quantify PI3P levels rely on radionuclide labelling techniques (Chicanne et al., 2012). We optimised a previously used label-free LC-MS/MS based method to quantify PI3P levels from *Drosophila* larval lysates [Fig 3B depicts a chromatogram derived from injecting wild type deacylated lipid samples] that allows the chromatographic separation and quantification of PI3P levels (Kiefer et al., 2010). To test if the ability of Mtm to dephosphorylate PI3P might be linked to its ability to reverse cell size in *dPIP4K^29^*, we measured PI3P levels in these genotypes. We observed that PI3P was significantly reduced in *dPIP4K^29^* larvae expressing Mtm compared to *dPIP4K^29^* (Figure 3C). We also measured PI3P levels from larvae expressing Mtm in an otherwise wild type background (Figure 3D) and found a modest reduction in the levels of PI3P. These results highlight the potential for PI3P levels to be correlated to the phenotype of cell size regulation in *Drosophila* salivary glands.

### dPIP4K regulates PI3P levels *in vivo*

Since reducing PI3P levels was correlated with cell size reversal in *dPIP4K^29^* (Figure 2A and 3C), we measured PI3P levels in *dPIP4K^29^*. Interestingly, we observed that PI3P was elevated in *dPIP4K^29^* larvae as compared to controls (Figure 4A). In order to confirm this observation of elevated PI3P levels in *dPIP4K*^29^ by an independent method, we devised an alternate assay to measure PI3P from larvae. Briefly, we developed an *in vitro* lipid kinase reaction using purified mCherry::dFab1 to quantify PI3P from larval lipid extracts using radionuclide labelling (schematic in Figure S4A). Figure 4B indicates the PI(3,5)P_2_ spot on a TLC formed from PI3P during the *in vitro* kinase reaction. Although lipid extracts from wild type and *dPIP4K^29^* larvae showed similar PI(3,5)P_2_ spot intensities on the TLC (Figure 4B), normalisation of the PI(3,5)P_2_ spot intensity against total organic phosphate levels in each sample confirmed that the total PI3P levels were higher in *dPIP4K^29^* (Figure 4C) compared to controls. To confirm that the increase of PI3P in *dPIP4K^29^* was a result of the absence of PIP4K, we reconstituted wild type dPIP4K in *dPIP4K^29^* and measured PI3P (*Act5C> dPIP4KeGFP; dPIP4K^29^*) and observed that the elevated PI3P in *dPIP4K^29^* was reverted to normal, indicating that dPIP4K can indeed regulate PI3P levels *in vivo* (Figure 4D). The catalytic activity of dPIP4K is essential to maintain salivary gland cell size (Mathre et al., 2019). Therefore, to check whether this catalytic activity was also necessary to control PI3P levels *in vivo*, we reconstituted *dPIP4K^29^* with a catalytically inactive dPIP4K (dPIP4K^D271A^) and measured PI3P (*Act5C> dPIP4K^D271A^; dPIP4K^29^*); we found that expressing catalytically dead dPIP4K^D271A^ could not significantly decrease the levels of PI3P in *dPIP4K^29^* (Figure 4E). These findings indicate that the catalytic activity of dPIP4K is required to regulate PI3P levels *in vivo*.

**Figure 4:**
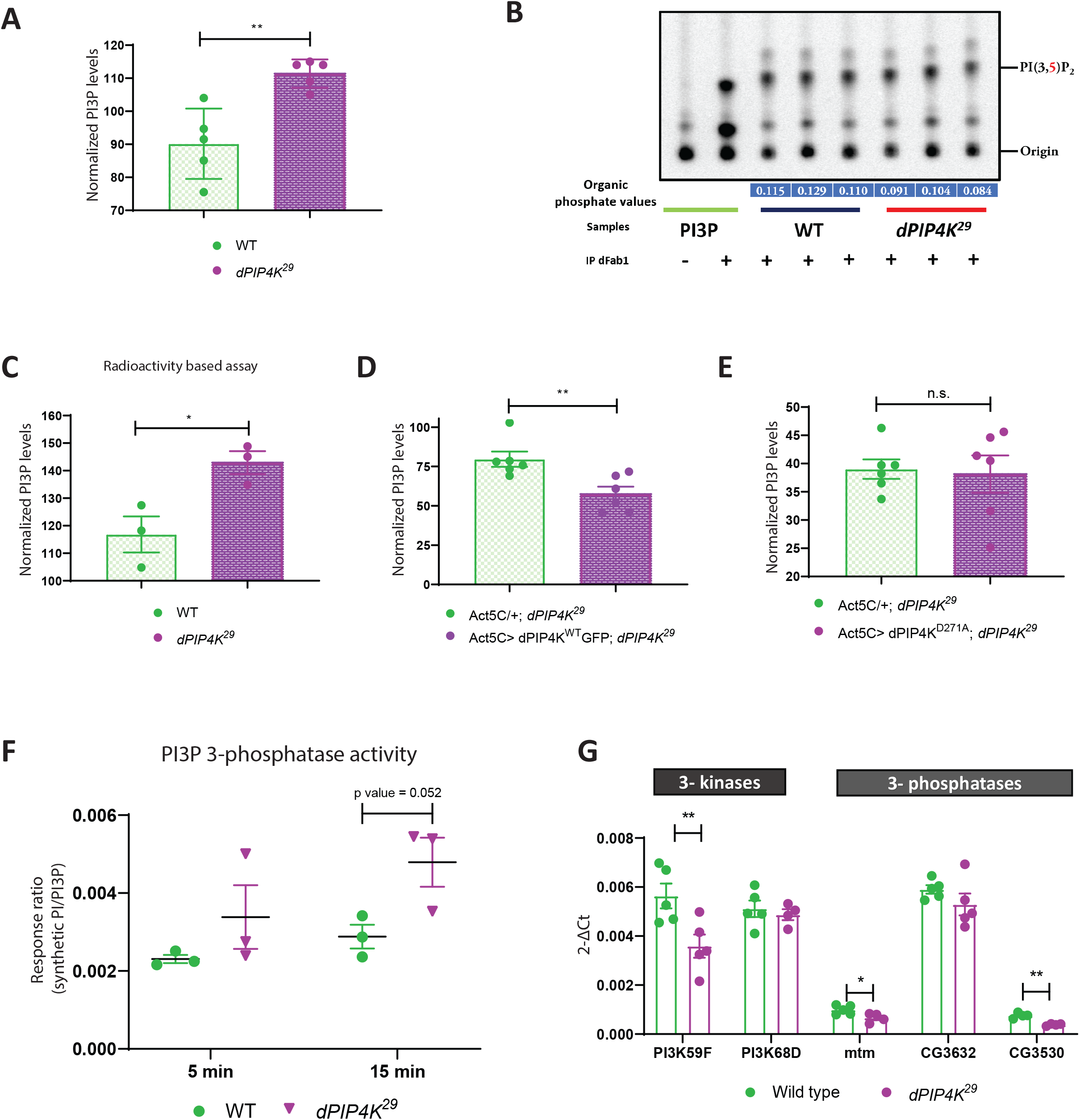
dPIP4K regulates PI3P levels *in vivo*. (A) Graph representing Normalised PI3P levels which is the peak area of GroPI3P/ peak area of GroPI4P normalised to organic phosphate value as mean ± S.E.M. of wild type (green) and *dPIP4K^29^* (majenta). Biological samples = 5, where each sample was made from five third instar wandering larvae. Unpaired t-test with Welch correction showed p value = 0.008. (B) Autoradiograph of TLC ran with lipid samples from *in vitro* PI3P mass assay using wild type (WT) and *dPIP4K^29^* lipid samples. The first two lanes from the left are obtained from mass assay reactions using synthetic PI3P standard without or with addition of dFab1 enzyme, respectively. The origin spot and PI(3,5)P_2_ spots are marked. (C) The graph represents normalised PI3P levels. Briefly, the spot marked as PI(3,5)P_2_ on TLC in (B) is obtained by converting PI3P in the samples using immunoprecipitated dFab1 in presence of ϒ^32^P-ATP are quantified using image analysis and then normalised to organic phosphate value (indicated in blue embedded text under TLC) to obtain normalised PI3P levels. Biological samples = 3, where each sample was made from five third instar wandering larvae. Student’s unpaired t-test with Welch correction showed p value = 0.036. (D) Graph representing Normalised PI3P levels which is the peak area of GroPI3P/ peak area of GroPI4P normalised to organic phosphate value of total lipid extracts as mean ± S.E.M. of Act5C/+; *dPIP4K^29^* (green), or Act5C> dPIP4K^WT^GFP, *dPIP4K^29^* (majenta). Biological samples = 5, where each sample was made from three third instar wandering larvae. Student’s unpaired t-test with Welch correction showed p value = 0.008. (E) Graph representing Normalised PI3P levels which is the peak area of GroPI3P/ peak area of GroPI4P normalised to organic phosphate value of total lipid extracts as mean ± S.E.M. of Act5C/+; *dPIP4K^29^* (green) or Act5C> dPIP4K^D271A^, *dPIP4K^29^* (majenta). Biological samples = 6, where each sample was made from three third instar wandering larvae. Student’s unpaired t-test with Welch correction showed p value = 0.818. (F) *In vitro* phosphatase assay on synthetic PI3P. Graph representing the response ratio of 17:0 20:4 PI (Product)/17:0 20:4 PI3P (Substrate) formed as mean ± S.E.M. on addition of either wildtype (WT) or *dPIP4K^29^* lysates for either a 5 min or a 15 min reaction. Lysate samples = 3, where each sample was made from five third instar wandering larvae. Multiple unpaired t-test showed p value = 0.26 for 5 min time point and p value = 0.052. (G) qPCR measurements for mRNA levels of *PI3K59F* and *PI3K68D* from either Wild type (green) or *dPIP4K^29^* (majenta). Student’s unpaired t-test showed p value = 0.01 for *PI3K59F* and p value = 0.58 for *PI3K68D*. qPCR measurements for mRNA levels of *Mtm, CG3632* and *CG3530* from either Wild type (green) or *dPIP4K^29^* (majenta). Student’s unpaired t-test showed p value = 0.03 for *Mtm*, p value = 0.23 for *CG3632* and p value = 0.0006 for *CG3530*.

### Regulation of PI3P by dPIP4K is unlikely to be via indirect mechanisms

Although PI5P is the canonical *in vivo* substrate for dPIP4K, the enzyme can also use PI3P as a substrate with low efficiency (Ghosh et al., 2019; Gupta et al., 2013), a feature conserved with mammalian PIP4Ks (Zhang et al., 1997). In the context of our observation that PI3P levels are elevated in *dPIP4K^29^*, dPIP4K could regulate PI3P levels either through its ability to directly phosphorylate this lipid or indirectly via its ability to regulate other enzymes that are established regulators of PI3P levels [e.g through negative regulation of PI 3-kinase activity or through positive regulation of a 3-phosphatase that dephosphorylate PI3P (Figure S4B)]. A reduction in 3-phosphatase activity on PI3P in *dPIP4K^29^* could lead to accumulation of PI3P. To test this possibility, the total 3-phosphatase activity of *dPIP4K^29^* lysates was assessed. We did not observe a reduction in 3-phosphatase activity that might explain the elevated PI3P levels. In fact, there was an increase in response ratio (indicative of the 3-phosphatase activity on PI3P) in a 15 minutes *in vitro* assay in mutant lysates as compared with wild type lysates (Figure 4F). In addition, we measured transcript levels of three putative 3-phosphatases – *Mtm*, *CG3632* and *CG3530* and found that the transcript levels of all the 3-phosphatases were unchanged in *dPIP4K^29^* as compared to controls, although there was an overall trend of decrease in all the genes (Figure 4G). PI3K59F activity could not be directly measured from larval lysates; however, we measured the mRNA expression of the two known PI 3-kinase genes – *PI3K59F* and *PI3K68D.* Although the transcript levels of *PI3K68D* was unchanged between *dPIP4K^29^* and controls, we observed that *PI3K59F* transcripts were in fact lower in *dPIP4K^29^* compared to controls (Figure 4G). Thus, it seems unlikely that upregulation of the aforementioned PI 3-kinases or downregulation of the 3-phosphatases contributes to the increased PI3P levels in *dPIP4K^29^*. These findings led us to conclude that the regulation of PI3P levels by dPIP4K is unlikely via indirect mechanisms.

### PIP4K regulates a non-endosomal PI3P pool in *Drosophila* salivary glands

The major source of PI3P generation in cells is the class III PI3-kinase called Vps34, whose *Drosophila* ortholog is PI3K59F. PI3K59F is known to be functional at two locations in cells – namely the early endosomal compartment and at multiple steps of autophagy pathway (Nascimbeni et al., 2017). In order to understand the location at which PI3P is elevated in *dPIP4K^29^*, we decided to restrict PI3K59F activity to revert the increased PI3P levels at both locations of PI3K59F activity. Consequently, if PI3P at either or both of these locations were relevant in regulating the cell size phenotype, we would achieve a reversal of cell size by down-regulating PI3K59F in *dPIP4K^29^* background. We down-regulated PI3K59F activity using RNA interference (Figure S5A depicts the extent of *PI3K59F* transcript knockdown) in *dPIP4K^29^* background and indeed observed a reversal of cell size (Figure 5A); knockdown of *PI3K59F* in an otherwise wild type background did not change cell size (Figure S5B). Further, measurement of PI3P from *dPIP4K^29^* larvae expressing PI3K59F RNAi showed a significant decrease in PI3P as compared to *dPIP4K^29^* larvae (Figure 5B).

**Figure 5:**
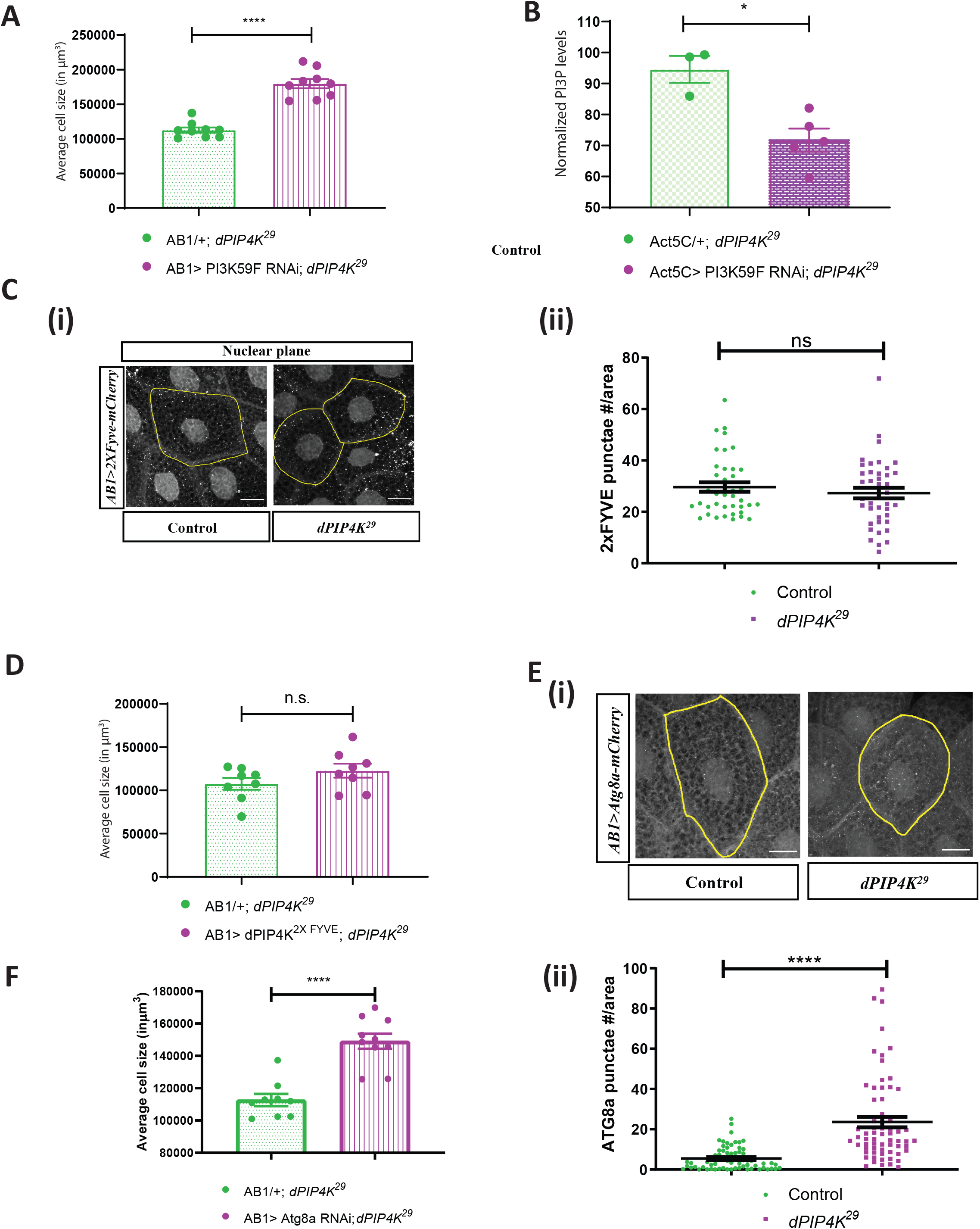
PIP4K in *Drosophila* salivary glands affects bulk autophagy to affect cell size. (A) Graph representing average cell size measurement (in μm^3^) as mean ± S.E.M. of salivary glands from wandering third instar larvae of *AB1/+ ; dPIP4K^29^* (n = 9), *AB1>PI3K59F* RNAi *; dPIP4K^29^* (n = 9). Sample size is represented on individual bars. Student’s unpaired t-test with Welch correction showed p value <0.0001. (B) Graph representing Normalised PI3P levels which is the peak area of GroPI3P/ peak area of GroPI4P normalised to organic phosphate value of total lipid extracts as mean ± S.E.M. of Act5C/+; *dPIP4K^29^* (green), or Act5C> dPIP4K^WT^GFP, *dPIP4K^29^* (majenta). Biological samples = 5, where each sample was made from five third instar wandering larvae. Student’s unpaired t-test with Welch correction showed p value = 0.011. (C) (i) Representative confocal z-projections depicting a sub population of early endosomal compartment using 2xFYVE-mCherry in the salivary glands from wandering third instar larvae of *AB1>* 2xFYVE-mCherry and *AB1>* 2xFYVE-mCherry *; dPIP4K^29^*. Scale bar indicated at 20 μm. (ii) Graph representing 2xFYVE punctae measurement in the salivary glands from wandering third instar larvae of *AB1>* 2xFYVEmCherry (N = 8, n=40) and *AB1>* 2xFYVEmCherry*; dPIP4K^29^* (N =8, n = 40). Student’s unpaired t-test with Welch correction showed p value = 0.4057 (D) Graph representing average cell size measurement (in μm^3^) as mean ± S.E.M. of salivary glands from wandering third instar larvae of *AB1/+ ; dPIP4K^29^* (n = 8), *AB1>dPIP4K^2XFYVE^; dPIP4K^29^* (n = 8). Sample size is represented on individual bars. Student’s unpaired t-test with Welch correction showed p value = 0.171. (E) (i) Representative confocal z-projections depicting autophagosomal levels using Atg8a-mCherry in the salivary glands from the wandering third instar larvae of *AB1>ATG8a-mCherry* and *AB1>ATG8a-mCherry; dPIP4K^29^*. Scale bar indicated at 20 μm. (ii) Graph representing Atg8a punctae measurement in the salivary glands from wandering third instar larvae of *AB1>ATG8a-mCherry* (N = 10, n = 60) and *AB1>ATG8a-mCherry; dPIP4K^29^* (N = 10, n = 62). Student’s unpaired t-test with Welch correction showed p value <0.0001. (F) Graph representing average cell size measurement (in μm^3^) as mean ± S.E.M. of salivary glands from wandering third instar larvae of *AB1/+ ; dPIP4K^29^* (n = 9), *AB1>PI3K59F* RNAi *; dPIP4K^29^* (n = 9). Sample size is represented on individual bars. Student’s unpaired t-test with Welch correction showed p value <0.0001.

To test if the early endosomal PI3P pool contributes to the reduced cell size in *dPIP4K^29^*, we imaged the tandem FYVE domain fused to mCherry (mCherry-2XFYVE), a reporter for endosomal PI3P, in salivary glands of *dPIP4K^29^* and compared it to wild type. The mCherry-2XFYVE probe revealed punctate structures which were perinuclear (Figure 5Ci). Quantification of the number of punctae per unit area calculated for the perinuclear sub-population showed no significant difference between wild type and *dPIP4K^29^* salivary glands (Figure 5Cii) although the probe was expressed at equal levels in both genotypes (Figure S5C). To further validate if change of PI3P at the endosomal location in *dPIP4K^29^* was correlated to the requirement of dPIP4K to support cell size, we tagged dPIP4K with the tandem FYVE domain at the C-terminus end of the protein (dPIP4K^2XFYVE^) to target it to the PI3P enriched endosomal compartment and reconstituted this in the background of *dPIP4K^29^*. We did not observe a significant change in the cell size of *dPIP4K^29^* under these conditions suggesting that dPIP4K function is dispensable at this location (Figure 5D) for cell size regulation.

The other sub-cellular location at which a Vps34-regulated PI3P pool is important, is the early autophagosomal membranes. We were unable to directly measure the PI3P pool at autophagosomal membranes. However, an increase in PI3P levels at this compartment would lead to an increase in the extent of autophagy (Burman and Ktistakis, 2010) and can be assayed by an increase in the *Drosophila* ortholog of microtubule-associated protein 1A/1B-light chain 3 (LC3) called Atg8a. We expressed mCherry::Atg8a in salivary glands and the probe was expressed at equal levels in both genotypes (Figure S5D). Measurement of the number of mCherry::Atg8a punctae showed a significant increase in *dPIP4K^29^* glands compared to controls (Figure 5Ei-ii), suggesting that the PI3P pool associated with the autophagy compartment is upregulated in *dPIP4K^29^*.

### PIP4K in *Drosophila* salivary glands affects bulk autophagy to affect cell size

PI3P is formed by Vps34 and regulated by Atg1 mediated activation of the Vps34 Complex I components Beclin-1 and Atg14, following which, lipidated Atg8a fuses with the formed omegasome membrane containing PI3P to mature into autophagosomes (King et al., 2021). It has been demonstrated that induction of autophagy by over-expressing Atg1 can cause a decrease in cell size of fat body cells in *Drosophila* larvae (Scott et al., 2007). Likewise, in this study, we found that the over-expression of Atg1 in the salivary glands of *Drosophila* larvae caused a drastic decrease in cell size (Figure S5E). Importantly, downregulating Atg1 activity in *dPIP4K^29^* could reverse the cell size phenotype in salivary glands while no change was observed in otherwise wild type background, indicating that the autophagy pathway is upregulated in *dPIP4K^29^* (Figure S5Fi-ii). We reasoned that if the elevated PI3P in *dPIP4K^29^* causes an upregulation in autophagy leading to cell size reduction, then by down-regulating Atg8a, we would be able to reverse the phenotype of cell size decrease. We indeed observed that down-regulation of Atg8a in *dPIP4K^29^* caused a reversal of the reduced cell size (Figure 5F), while there was no significant change in cell size by down-regulation of Atg8a in otherwise wild type background (Figure S5G). Further, down-regulation of Atg8a using the same RNAi line was able to decrease the number of mCherry::Atg8a in salivary glands (Figure S5Hi-ii); the expression of the probe being equivalent between both genotypes (Figure S5I).

### PI3P regulates cell size in salivary glands

We tested the effect of modulating PI3P levels in otherwise wild type salivary glands. Depletion of *Mtm*, using an RNAi line has previously shown to increase PI3P levels in *Drosophila* (Jean et al., 2012; Velichkova et al., 2010). Using qPCR analysis, we validated that the RNAi reagent causes specific down-regulation of *Mtm* transcripts in *Drosophila* (Figure 6A). Interestingly, expressing Mtm RNAi in salivary glands caused a significant decrease in cell size (Figure 6Bi and ii). Myotubularins are known to dimerize in cells and Mtm harbours a C-terminal coiled-coil domain which can potentially aid in dimerization (Jean et al., 2012). We observed a similar but smaller reduction in cell size when a catalytically dead version of Mtm (Mtm^D402A^), that is expected to act as dominant negative construct was expressed in salivary glands (Figure S6A). Further, measurement of PI3P levels revealed a modest upregulation of PI3P levels when measured from larvae expressing the Mtm RNAi (*da*> *Mtm RNAi*, Figure 6C). Therefore, Mtm inhibition in an otherwise wildtype background can cause PI3P elevation and cell size decrease in *Drosophila*.

**Figure 6:**
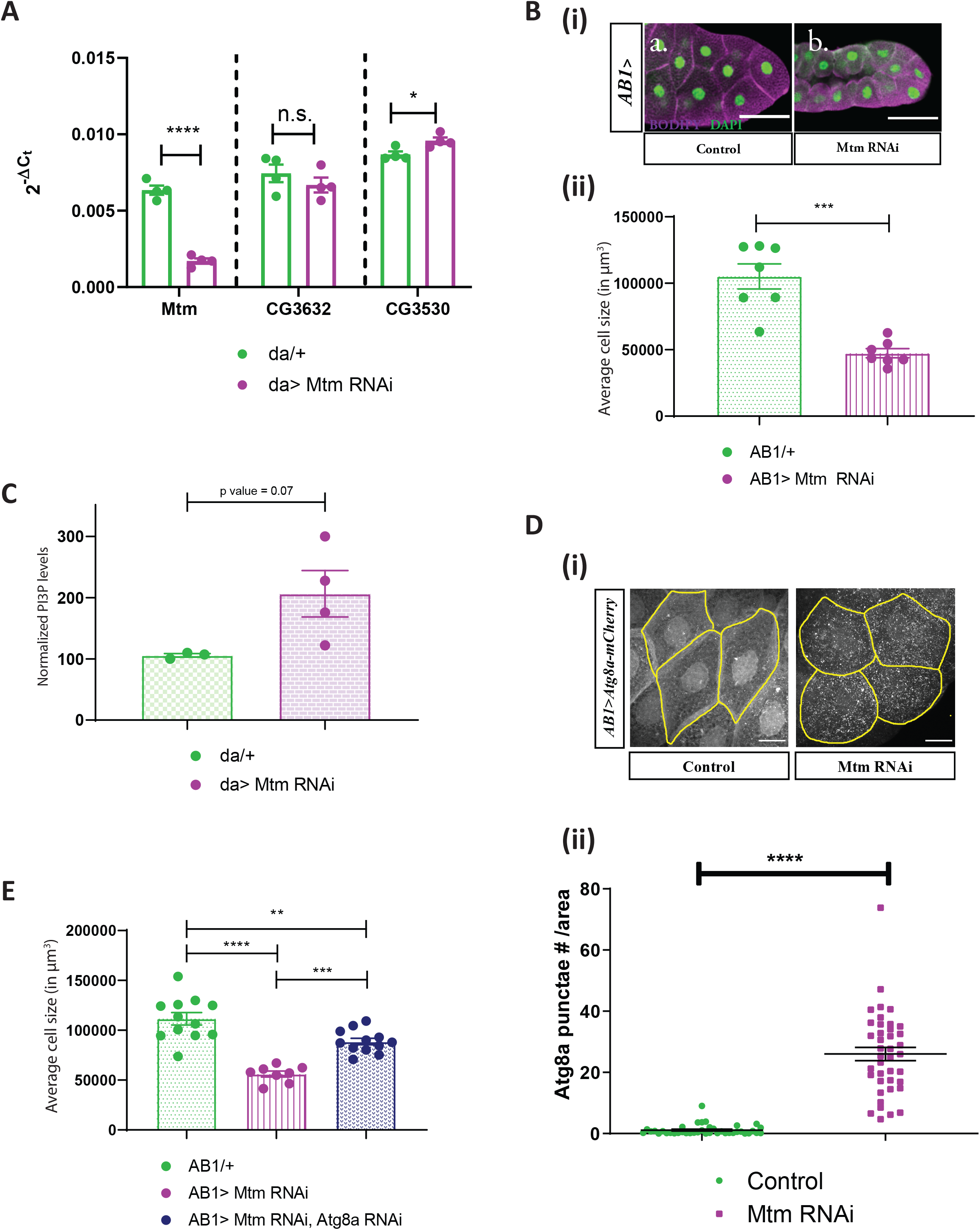
PI3P regulates cell size in salivary glands. (A) qPCR measurements for mRNA levels of *mtm, CG3632 and CG3530* from either Control (*daGal4*/+, in green) or *da*> *Mtm* RNAi, in majenta. Multiple t-test with post hoc Holm-Sidak’s test showed p value < 0.0001 between *daGal4*/+ and *da*> *Mtm* RNAi for *Mtm* and p value = 0.35 between *daGal4*/+ and *da*> *Mtm* RNAi for *CG3632*, and p value = 0.04 between *daGal4*/+ and *da*> *Mtm* RNAi for *CG3530*. (B) (i) Representative confocal images of salivary glands from the genotypes a. *AB1Gal4/+,* b. *AB1>Mtm* RNAi. Cell body is marked majenta by BODIPY conjugated lipid dye, nucleus is marked by DAPI shown in green. Scale bar indicated at 50 μm. (ii) Graph representing average cell size measurement (in μm^3^) as mean ± S.E.M. of salivary glands from wandering third instar larvae of *AB1Gal4/+* (n = 7), *AB1> Mtm* RNAi (n = 7). Sample size is represented on individual bars. Student’s unpaired t-test with Welch correction showed p value = 0.0005. (C) Graph representing Normalised PI3P levels which is the peak area of GroPI3P/ peak area of GroPI4P normalised to organic phosphate value as mean ± S.E.M. of *da*/+ (green) and *da> Mtm* RNAi (majenta). Biological samples = 4, where each sample was made from five third instar wandering larvae. Student’s unpaired t-test with Welch correction showed p value = 0.07. (D) (i) Graph representing Atg8a punctae measurement in the salivary glands from wandering third instar larvae of *AB1>ATG8a-mCherry* (N =7, n =40), *AB1>ATG8a-mCherry ; Mtm RNAi* (N =7, n =40). Student’s unpaired t-test with Welch correction showed p value <0.0001. (ii) Representative confocal z-projections depicting autophagosomal levels using Atg8a-mCherry in the salivary glands from the genotypes a. *AB1>ATG8a-mCherry,* b. *AB1>ATG8a-mCherry; Mtm RNAi.* Scale bar indicated at 20 μm. (E) Graph representing average cell size measurement (in μm^3^) as mean ± S.E.M. of salivary glands from wandering third instar larvae of *AB1/+* (n = 11), *AB1>Mtm* RNAi (n = 8), *AB1>Mtm* RNAi, *Atg8a* RNAi (n = 12). Sample size is represented on individual bars. One way ANOVA with post hoc Tukey’s test showed p value < 0.0001 between *AB1/+* and *AB1>Mtm* RNAi and p value = 0.0002 between *AB1/+ ; dPIP4K^29^* and *AB1>Mtm* RNAi, *Atg8a* RNAi.

To understand if the reduction in cell size brought about by down-regulating Mtm in salivary glands causes an upregulation of the autophagic pathway much like in *dPIP4K^29^* mutants, we measured the number of mCherry::Atg8a in salivary glands of Mtm RNAi (Figure 6Di). It was observed that there was a substantial increase in the number of mCherry::Atg8a punctae as quantified in Figure 6Dii, although the expression of the probe was expressed equivalent in both genotypes (Figure S6B). Interestingly, depleting Atg8a in salivary glands also depleted of Mtm could partially rescue cell size as compared to glands where Mtm alone was down-regulated (Figure 6E). These findings corroborate the relationship of increased PI3P levels to the upregulation of autophagy which can eventually contribute to a decrease in cell size of salivary glands of *Drosophila* larvae.

## Discussion

Conceptually, the cellular function of any enzyme can be considered to arise from its ability to regulate the levels of either the substrate or product. When PIP4K was originally described (Rameh et al., 1997), its ability to generate the product PI(4,5)P_2_ was recognised. However, PI(4,5)P_2_, can also be synthesized from phosphatidylinositol 4 phosphate (PI4P) by the activity of phosphatidylinositol 4 phosphate 5 kinase (PIP5K) [reviewed in (Kolay et al., 2016)]. Since PI4P is ca.10 times more abundant than PI5P, loss of PIP4K activity is unlikely to impact the overall levels of cellular PI(4,5)P_2_. Consistent with this, knockout of PIP4K does not reduce the overall level of PI(4,5)P_2_ [(Gupta et al., 2013) discussed in (Kolay et al., 2016)]. Further, a switch mutant version of PIP4K that can generate PI(4,5)P_2_ from PI4P but not PI5P, was unable to rescue the reduced cell size in *dPIP4K^29^* implying that the biochemical basis of dPIP4K function in supporting cell size is not its product PI(4,5)P_2_. Rather, given that the kinase activity of the enzyme is required for normal salivary gland cell size (Mathre et al., 2019), our findings imply that the levels of the substrate are likely to be relevant. It was vital to be able to measure the levels of putative substrates of PIP4K from *Drosophila* tissues and therefore, we developed a label-free LC-MS/MS based methods to detect and quantify PI3P and PI5P. Previously researchers used more cumbersome radioactivity based detection to measure PI3P and PI5P (Chicanne et al., 2012; Jones et al., 2013), however, new label-free methods are being reported to quantify PIPs (Ghosh et al., 2019; Morioka et al., 2022). In this study, we report the use of a label-free LC-MS/MS based method to measure PI3P levels *in vivo* from *Drosophila* tissues for the first time. In future, this method can also be used to measure PI3P levels from tissues of other model organisms to address key questions in PI3P biology.

Given that previous studies have identified PI5P as the substrate best utilised by PIP4K and that PI5P levels are elevated in *dPIP4K^29^,* we expected that the cell size phenotype will be mediated by PI5P levels. In the course of this study, we found that (i) the expression of Mtm, a 3-phosphatase that is able to generate PI5P *in vitro* from PI(3,5)P_2_ rescued the reduced cell size in *dPIP4K^29^*, (ii) The rescue of cell size in *dPIP4K^29^* by Mtm overexpression was not associated with a change in the levels of PI5P. These observations are not consistent with a role for elevated PI5P levels in the reduced cell size phenotype of *dPIP4K^29^*. Since PI3P has also been shown to be a substrate of dPIP4K, albeit with less efficiency (Ghosh et al., 2019; Gupta et al., 2013), we investigated PI3P levels and found that (i) PI3P levels were elevated in *dPIP4K^29^*, (ii) were reverted to wild type by reconstitution with a wild type dPIP4K transgene but not a kinase dead version, (iii) the rescue of cell size in *dPIP4K^29^* by expression of Mtm was associated with a reduction in the elevated PI3P levels. Together, these findings strongly suggest that the elevated PI3P levels in *dPIP4K^29^* underpin the reduced salivary gland cell size. If elevated PI3P is a regulator of cell size, then elevation of PI3P levels in wild type cells might also result in reduced cell size. Our observation that depletion of Mtm in a wildtype background results in elevated PI3P levels and also reduced cell size supports this model. Overall, our data supports a role for dPIP4K in the regulation of PI3P levels and cell size.

Interestingly, a recent study in MEFs grown in culture and downregulated of PIP4Kγ activity showed an increase in PI3P and PI(3,5)P_2_ levels along with an expected rise in PI5P levels (Al-Ramahi et al., 2017). In mammals, the major route of synthesis of PI3P is through the action of a class III PI3-kinase called Vps34. We observed that down-regulating PI3K59F, the ortholog of Vps34 in *Drosophila* reversed cell size of *dPIP4K^29^* salivary glands. These findings also identify PIP4K as a new regulator of PI3P levels along with Vps34.

In spite of having specific stereo-chemistries, there are instances which demonstrate PI3P and PI5P to be very similar to each other. The <u>F</u>ab1 (yeast orthologue of PIKfyve), <u>Y</u>OTB, <u>V</u>ac 1 (vesicle transport protein), and <u>E</u>EA1 (FYVE) domain has been used extensively for its lipid binding affinity for PI3P (Gillooly et al., 2001). But NMR analysis revealed that FYVE domain show lesser albeit significant binding affinity towards PI5P (Kutateladze et al., 1999). The <u>P</u>lant-<u>H</u>omeo-<u>D</u>omain (PHD) of ING2 protein, which has been used in quite a few studies to probe PI5P location have revealed secondary avidities for PI3P (Gozani et al., 2003). However, PIP4Kα has a less but significant *in vitro* kinase activity measured with PI3P as substrate (Ghosh et al., 2019; Zhang et al., 1997). Consequently, purified *Drosophila* PIP4K also shows a faint PI(3,4)P_2_ spot measured through radioactive kinase assay indicating its 4-kinase activity on PI3P (Gupta et al., 2013). However, due to the fold difference in *in vitro* activity between the two substrates, it was never envisioned that PIP4K could regulate PI3P levels *in vivo*. Our results indicate a possibility where the PIP4K can access a pool of PI3P *in vivo*, such that the enzyme achieves its optimal conditions for a successful kinase reaction to metabolise PI3P.

What is the mechanism by which cell size is reduced in *dPIP4K^29^*? It has been reported in several studies in mammalian models that loss of PIP4K function is associated with an increase in either the initiation step or flux of autophagy (Al-Ramahi et al., 2017; Lundquist et al., 2018; Vicinanza et al., 2015). Moreover, it has been shown in human cells that PI5P can initiate autophagy and can even take over the function of PI3P to initiate autophagy in wortmannin-treated cells (Vicinanza et al., 2015). We found that just by altering PI3P levels without any change in PI5P levels, we could modify the phenotype of cell size of *dPIP4K^29^*. PI3P has been reported in primarily two cellular compartments, early endosomes and autophagosomes. TORC1 activity is reported to be downregulated in *dPIP4K^29^* (Gupta et al., 2013) and consistent with the function of TORC1 in regulating autophagy, we found that levels of autophagy in *dPIP4K^29^* salivary gland cells was increased. Reducing PI3P levels by genetic knockdown of Vps34 (Class III PI3K) reversed the reduced cell size phenotype in *dPIP4K^29^* implying a role for Vps34 synthesized PI3P in regulating cell size. Further, an early endosome specific dPIP4K construct could not revert the cell size change of *dPIP4K^29^*, indicating that the relevant pool of PI3P that regulates cell size is not at the early endosome. Finally, inhibition of autophagy by down-regulation of Atg8a, a protein required downstream of PI3P formation at the phagophore membrane during autophagosome biogenesis, in wild type results in reduced cell size underscoring the requirement of normal levels of autophagy in controlling cell size in the salivary gland. Additionally, we show that *Drosophila* Mtm also regulates cell size by downmodulating PI3P levels and autophagy. Recent studies in *Drosophila* have identified a role for *CG3530/Mtmr6* in control of basal autophagy in fat bodies (Allen et al., 2020; Manzéger et al., 2021). However, we do acknowledge that Mtm downregulation does affect the endosomal PI3P pool as is reported earlier from the Kiger lab (Velichkova et al., 2010). With the present data we cannot rule out an effect of endosomal PI3P in contributing to cell size regulation in Mtm downregulation and perhaps the partial rescue of cell size by down-regulating Atg8a explains this phenomenon (Figure 6E). Together our data provide compelling evidence that the elevated levels of PI3P in *dPIP4K^29^* induces enhanced autophagy leading to reduction in cell size.

## Materials and methods

### Fly strains and stocks

All experiments were performed with Drosophila melanogaster (hereafter referred to as Drosophila). Cultures were reared on standard medium containing corn flour, sugar, yeast powder and agar along with antibacterial and antifungal agents. Genetic crosses were set up with Gal4 background strains and maintained at 25°C and 50% relative humidity (Brand and Perrimon, 1993). There was no internal illumination within the incubator and the larvae of the correct genotype was selected at the 3^rd^ instar wandering stage using morphological criteria. Drosophila strains used were Oregon-R and w^1118^ (wild type strain), dPIP4K^29^ (homozygous null mutant of dPIP4K made by Raghu lab), da-Gal4, Act5C-Gal4/CyoGFP, AB1-Gal4, UAS hPIP4K2B/TM6Tb, UAS hPIP4K2B^[A381E]^/TM6Tb, Mtm^WT^GFP (Amy Kiger, UCSD), Mtm-IR (#AK0246, Amy Kiger, UCSD), UAS dPIP4K^WT^eGFP, UAS dPIP4K^[D271A]^(untagged), UAS PI3K59F RNAi (v100296, VDRC), Atg1 RNAi (44034, Bloomington), Atg8a RNAi (34340, Bloomington), w; UAS-mCherry:2XFYVE^2^ (Amy Kiger, UCSD), UAS-mCherry-Atg8a (37750, Bloomington).

### S2R+R+ cells: culturing and transfection

Drosophila S2R+R+ cells were cultured and maintained as mentioned earlier (Gupta et al., 2013). Transient transfections for 48 hours were performed as mentioned previously (Mathre et al., 2019). Primers for amplifying dsRNA template against Drosophila genes were selected from DRSC/TRiP Functional Genomics Resources after confirming specificity of primers. A T7 RNA polymerase promoter sequence (5’-TAATACGACTCACTATAGGGAGA-3’) was added at the 5’ end of the primers for the T7 DNA dependent RNA polymerase to bind during in vitro transcription. The dsRNA was synthesised using amplicons amplified from BDGP gold clones (Mtm: LD28822, CG3632: LD11744 and CG3530: GH04637), purchased from DGRC. Following are the list of primers used for the in vitro transcription of dsRNA:

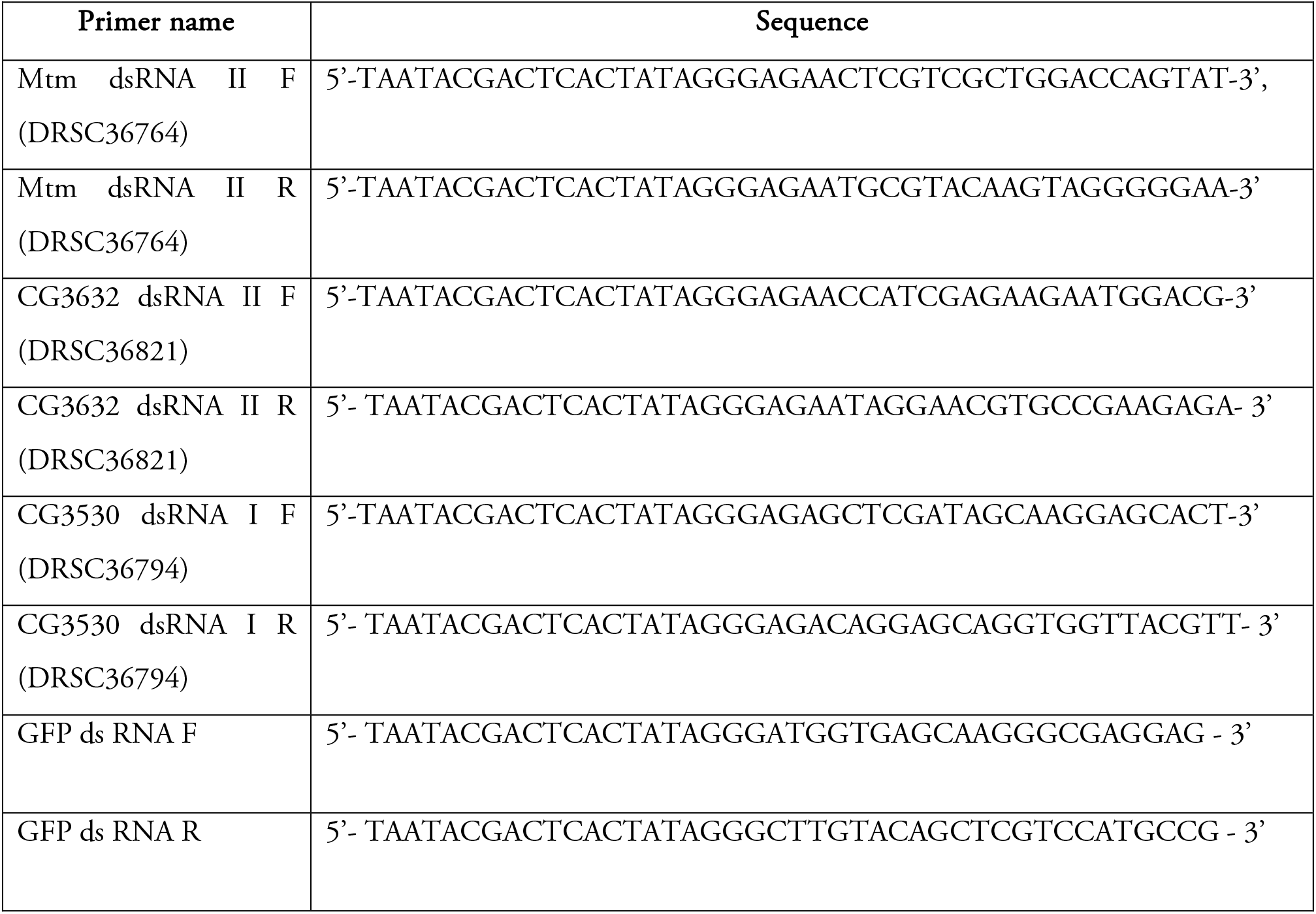

### RNA extraction and qPCR analysis

RNA was extracted from Drosophila S2R+R+ cells using TRIzol reagent (15596018, Life Technologies, California, USA). Purified RNA was treated with amplification grade DNase I (18068015, Thermo Fisher Scientific, California, USA). cDNA conversion was done using SuperScript II RNase H– Reverse Transcriptase (18064014, Thermo Fisher Scientific) and random hexamers (N8080127, Thermo Fisher Scientific). Quantitative PCR (Q-PCR) was performed using Power SybrGreen PCR master-mix (4367659, Applied Biosystems, Warrington, UK) in an Applied Biosystem 7500 Fast Real Time PCR instrument. Primers were designed at the exon-exon junctions following the parameters recommended for QPCR. Transcript levels of the ribosomal protein 49 (RP49) were used for normalization across samples. Three separate samples were collected from each treatment, and duplicate measures of each sample were conducted to ensure the consistency of the data. The primers used were as follows:

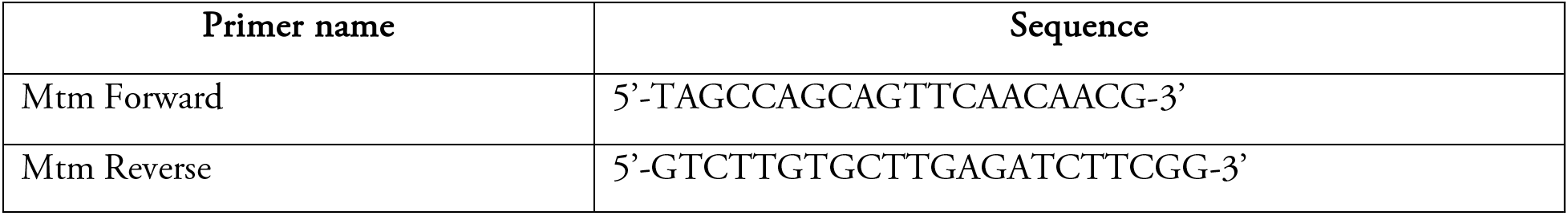
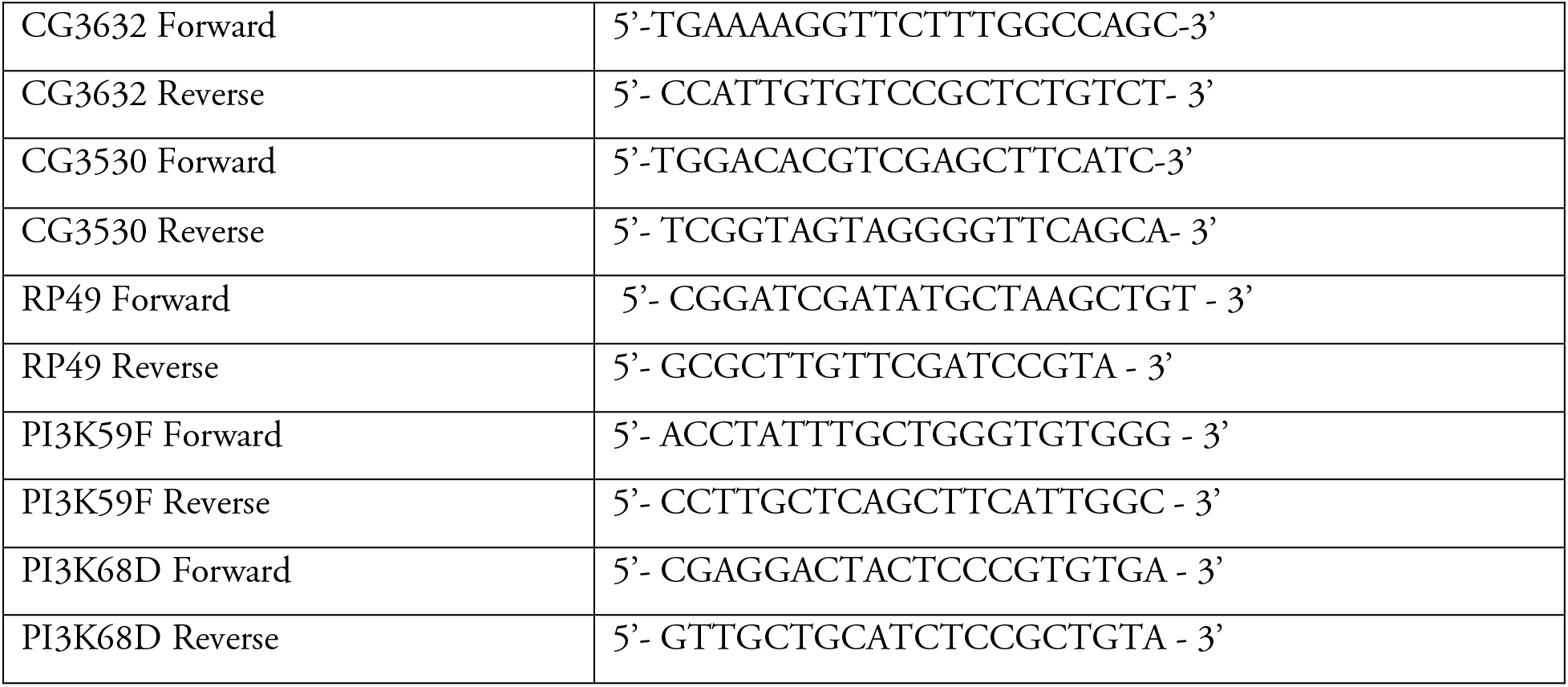

### Western blotting and immuno-precipitation

<u>Westerns:</u> Salivary glands or larval samples were made exactly as mentioned in our previous work (Ghosh et al., 2019; Mathre et al., 2019). Dilutions of antibodies used: 1:4000 for anti-tubulin (E7-c), (mouse) from DSHB, 1:1000 for anti-mCherry antibody (Cat# PA5-34974) (Rabbit) from Thermo, 1:1000 for anti-HA antibody (Cat# 2367S) (Mouse) from CST and Normal Rabbit IgG (sc-2027) from Santa Cruz. <u>Immuno-precipitation:</u> About 2 million S2R+R+ cells were transfected for 48 hours and lysates were prepared using 200 µl of same lysis buffer used for the preparation of protein samples for western blotting. After lysis for 15 mins at 4 °C, the samples were spun down at 13000 X g for 10 mins to remove cellular debris. 5% of the supernatant obtained was kept aside for input control; to the rest of the sample lysis buffer was added to make up the volume to 1 mL. The volumes were split in two halves – one for IgG control and the other for immune-precipitation. About 1.6 µg equivalent of antibody/normal rabbit IgG was used for over-night incubation at 4 °C with continuous rotation. mCherry tagged Drosophila Fab1 complexes with anti-mCherry antibody were precipitated by ∼60 µl slurry of washed and blocked protein-G sepharose beads (according to manufacturer’s protocol, Sigma # GE17-0886-01) for 2 hours at 4 °C. The beads were then washed with 0.1% TBST containing 0.1% 2-Mercaptoethanol, 0.1mM EGTA for two times and resuspended in 100 µl of the same buffer and stored at 4 °C till kinase assay was performed.

### Cell size measurements

Salivary glands were dissected from wandering third instar larvae and fixed in 4% paraformaldehyde for 20 min at 4°C. Post fixation, glands were washed thrice with 1X Phosphate Buffered Saline (PBS) and incubated in BODIPY™ FL C12-Sphingomyelin (Cat# D7711) for 3 hours at room temperature, following which they were washed thrice in 1X PBS and stained with either DAPI (Thermo Fisher, cat# D1306) or TOTO3 (Thermo Fisher, cat# T3604) for 10 mins at room temperature and washed with 1X PBS again. About 2-3 glands per slide were then mounted in 70% glycerol and imaged. Imaging was done on Olympus FV1000 or FV3000 Confocal microscope using a 10X objective. The images were then stitched into a 3D projection using an ImageJ plugin. These reconstituted 3D z stacks were then analyzed for nuclei numbers (for cell number) and volume of the whole gland using Volocity Software (version 5.5.1, Perkin Elmer Inc.). The average cell size was calculated as the ratio of the average volume of the gland to the number of nuclei.

### Atg8a punctae measurements

Around 40 first instar larvae were picked and incubated per vial to control for crowding. Salivary glands were dissected from wandering third instar larvae and fixed in 2.5% paraformaldehyde for 20 min at room temperature. Post fixation, glands were washed twice with 1X PBS. Glands were mounted in 70% glycerol and imaged on the same day. Imaging was done on an Olympus FV3000 Confocal microscope using a 60X objective. The 3D images were stitched to give one 2D image using Zproject in ImageJ. These 2D images were then analysed for the number of punctae using the 3D object counter plugin in ImageJ. The number of punctae were normalised to the area of the cell and plotted for the respective genotypes.

### 2xFYVE punctae measurements

Salivary glands were dissected and imaged as described for the ATG8a punctae measurements. The 3D images were stitched to give one 2D image using Zproject in ImageJ. The 2D images were then analysed for the number of punctae (analysed using 3D object counter plugin in ImageJ). The number of punctae were normalised to the area of the cell and plotted for the respective genotypes.

### Lipid standards

diC16-PI3P – Echelon P-3016; diC16-PI4P – Echelon P-4016; Avanti 850172 | rac-16:0 PI(3,5)P_2_-d5 (Custom synthesised) ; 17: 0 20: 4 PI3P - Avanti LM-1900 ; 17: 0 20: 4 ; PI(4,5)P_2_ - Avanti LM-1904.

### Radioactivity based PI3P mass assay

diC16-PI3P (Echelon) was mixed with 20 µM Phosphatidylserine (PS) (Sigma #P5660) and dried in a centrifugal vacuum concentrator. For biological samples, PS was added to the organic phase obtained at the end of the neomycin chromatography before drying. To the dried lipid extracts, 50 µl 10 mM Tris-HCl pH 7.4 and 50 µl diethyl ether was added and the mixture was sonicated for 2 mins in a bath sonicator to form lipid micelles. The tubes were centrifuged at 1000 X g to obtain a diethyl ether phase and vacuum centrifuged for 2 mins to evaporate out the diethyl ether. At this time, the reaction was incubated on ice for about 10 mins and 2X kinase assay buffer (100 mM Tris pH 7.4, 20 mM MgCl_2_, 140 mM KCl, and 2 mM EGTA and 10 μL immuno-precipitated dFab1 bead slurry was added. To this reaction, 10 μCi [γ-^32^P] ATP was added and incubated at 30 °C for 16 hours. Post 16 hours, the lipids were extracted from the reaction as described earlier in a radioactive PI5P mass assay protocol (Jones et al., 2013).

### Thin layer Chromatography

Extracted lipids were resuspended in chloroform and resolved by TLC (preactivated by heating at 90^0^C for 1 hour) with a running solvent (45:35:8:2 chloroform: methanol: water:25% ammonia). Plates were air dried and imaged on a Typhoon Variable Mode Imager (Amersham Biosciences).

### In Vitro dFab1 immunoprecipitate based Lipid 5-kinase assays

600 picomoles of either 17:0 | 14:1 PI (Avanti # LM 1504) or 17:0 | 20:4 PI3P (Avanti # LM 1900) were mixed with 20 µl of 0.5 (M) of Phosphatidylserine (PS) (P5660, Sigma) and dried in a centrifugal vacuum concentrator. To this, 50 µl 10 mM Tris-HCl pH 7.4 was added and the mixture was sonicated for 3 minutes in a bath sonicator to form lipid micelles. At this time, the reaction was incubated on ice for ∼ 10 minutes and 2X kinase assay buffer (100 mM Tris pH 7.4, 20 mM MgCl_2_, 140 mM KCl, and 2 mM EGTA and equal volumes of immunoprecipitated dFab1 was added. For LC-MS/MS based experiments the kinase assay buffer contained 80 μM cold ATP (10519979001, Roche)). The rest of the procedure was followed as mentioned in the following section.

### In Vitro larval lysate-based Lipid 3-phosphatase assays

The assay conditions have been adopted from a previous study (Schaletzky et al., 2003). The phosphatase assay comprises three parts-<u>(i) preparation of lysate:</u> third instar wandering larvae were collected in groups of 5 and lysed in phosphatase lysis buffer containing 20 mM Tris-HCl (pH 7.4), 150 mM NaCl, 1% Triton X-100 (v/v) and protease inhibitor cocktail (Roche), by incubating the resuspended mixture in ice for 15-20 min. The larval carcasses were removed by a brief spin for 5 mins at 1000 x g speed. Total protein was estimated by Bradford’s reagent and desired amount of lysate was used for the subsequent assay. <u>(ii) Lipid</u> <u>phosphatase assay:</u> 600 picomoles of either 17:0 20:4 PI3P or d5-PI(3,5)P_2_ lipid was mixed and dried with 20 µl of 0.5 M bovine brain derived Phosphatidylserine (PS) (Sigma #P5660) followed by bath sonication of the mixture in presence of 50 µl of 10 mM Tris-HCl (pH 7.4) for 3 min at maximum amplitude. To this 50 µl of 2× phosphatase assay buffer (Schaletzky et al., 2003) and 10 µg total protein equivalents of cell free lysate was added and the reaction was incubated for 15 min at 37 °C. The reaction was quenched with 125 µl of 2.4 (N) HCl followed by lipid extraction described earlier (Jones et al., 2013). Samples for the PI3P assay were processed according to section (iii). For the PI(3,5)P_2_ phosphatase assay the dried lipids from this step were resuspended in 20 µl of 0.5 M PS and dried. To this, 50 µl of 10 mM Tris-HCl (pH 7.4) was added and bath sonicated for 3 min similar to the first step of the assay. At this step, 50 µl of 2× kinase buffer containing 80 µM O^18^ ATP (OLM-7858-PK, Cambridge Isotope laboratories, Inc) and 1 µg of bacterial purified human PIP4Kα-GST was added and the reaction was incubated at 30 °C for 1 hour. This was followed by lipid extraction as described in the previous step. <u>(iii) Derivatization</u> <u>of lipids and LC-MS/MS:</u> The organic phases were collected from the last step and dried and 50 μl of 2M TMS-diazomethane (Acros #AC385330050) was added to each tube and vortexed gently for 10 min at room temperature. The reaction was neutralized using 10 μl of glacial acetic acid. The samples were dried in vacuo and 200 µl of methanol was used to reconstitute the sample to make it ready for injection for LC-MS/MS analysis.

### Lipid isolation for PI5P and PI3P measurements

Lipids from larvae were isolated from 3 or 5 third instar wandering larvae for PI5P or PI3P measurements, respectively. Total lipids were isolated and neomycin chromatography (for PI5P measurements only) was performed as described earlier (Ghosh et al., 2019).

### LC-MS/MS for in vitro assays and PI5P measurements

The instrument operation was followed similar to the description in our previous methods work on PI5P quantification (Ghosh et al., 2019). For in vivo lipid measurements, the samples were washed with post-derivatisation wash step before injecting in mass spec. Samples were run on a hybrid triple quadrupole mass spectrometer (Sciex 6500 Q-Trap or Sciex 5500 Q-Trap) connected to a Waters Acquity UPLC I class system. Separation was performed on a ACQUITY UPLC Protein BEH C4, 300Å, 1.7 µm, 1 mm X 100 mm column [Product #186005590] using a 45% to 100% acetonitrile in water (with 0.1% formic acid) gradient over 10 mins. MS/MS and LC conditions used were as described earlier (Ghosh et al., 2019).

### Larval PI3P measurements

We adopted a previously used method of deacylation of total lipids followed by detection by LC-MS/MS using ion-paring based separation chemistry followed by detection using mass spec (Jeschke et al., 2015; Kiefer et al., 2010). Using our conditions, we could not reproducibly separate the deacylated PI5P isomeric peak from biological samples. But we could always separate deacylated PI3P from PI4P in these biological samples (Figure 3B). Synthetic standards were used to determine the retention times of the individual peaks. Figure S3A shows synthetic GroPI3P at Rt = 6.13 min and GroPI4P at Rt = 6.95 min. The Rt of GroPI3P and GroPI4P was shifted in case of biological samples and in order to confirm the peaks were representative of the corresponding analytes, we spiked synthetic GroPI3P into the biological sample of Figure S3B. As expected, we observed a spike in the first peak, albeit at Rt = 7.65 min, without a significant change in the second peak at Rt = 8.70 min, thus confirming that the first peak was indeed corresponding to PI3P. Further, we also verified that GroPI3P can be linearly detected at a range of 30 - 3000 picograms on column and GroPI4P can be linearly detected at a range of 30 - 4000 picograms on column (Figure S3C and S3D). We determined that the Limit of detection (LOD) was 20 picograms on column for GroPI3P and GroPI4P as concluded from Signal to Noise (S/N) being 30 and 24, respectively.

The following are the steps by which PI3P measurements were performed: <u>(i) Larval lipid extraction:</u> As mentioned in previous section <u>(ii) lipid deacylation:</u> Dried lipid extracts were incubated with 1 mL of 25% methylamine solution in water/methanol/n-butanol (43:46:11) at 60 °C for 1 hour followed by drying this extract in vacuo (∼ 3-4 hours). <u>(iii) fatty acid wash:</u> Next, the lipids were reconstituted in 40-50 µl MS-grade water and to this an equal volume fatty acid extraction reagent (1-butanol/petroleum ether (40–60 °C boiling)/ethyl formate in a ratio of 20/4/1 (vol/vol/vol)) was added and vortexed for 2 mins. Following this, the tubes were centrifuged for 5 mins at 1000 x g to obtain phase separation. The upper organic phase was discarded, and the lower aqueous phase was processed for LC-MS/MS analysis.

MRM values for commercial lipids used:

| Lipids | Parent ion | Daughter ion |
| --- | --- | --- |
| d5-diC16-PI5P | 938.5 | 556.5 |
| d5-diC16-PI(3,5)P<sub>2</sub> | 1046.5 | 556.5 |
| d5-diC16-<sup>18</sup>O-PIP<sub>2</sub> | 1052.5 | 556.5 |
| 17:0\| 20:4 PI3P | 995.5 | 613.5 |
| 17:0\| 20:4 PIP<sub>2</sub> | 1103.5 | 613.5 |
| 17:0\|14:1 PI | 809.4 | 535.4 |
| 17:0\|14:1 PIP | 917.4 | 535.4 |

### LC-MS/MS for deacylated PI3P measurements

Deacylated PIPs (GroPI3P and GroPI4P) were run on a hybrid triple quadrupole mass spectrometer (Sciex 6500 Q-Trap) connected to a Waters Acquity UPLC I class system. Separation was performed on a Phenomenex Synergi™ 2.5 µm Fusion-RP 100 Å, LC Column 100 x 2 mm, [Product # 00D-4423-B0] maintained at 32°C during the run. Mobile phase A consisted of 4 mM DMHA and 5 mM acetic acid in water, and mobile phase B consisted of 4 mM DMHA and 5 mM acetic acid in 100% methanol. Flow rate was 0.2 mL/min.

The process of linear gradient elution was conducted as follows: 0–2 min (methanol, 3%), 2–5 min (methanol, 7%), 5–8 min (methanol, 12%), and 8–9 min (methanol, 100%). For next 4 min, solvent B was maintained at 100%. Then, equilibration was performed between 12.2 and 20.0 min using 3% methanol. The injection volume and running time of each sample was 3.0 µL and 20.0 min, respectively.

Mass spectrometry data was acquired in multiple reaction monitoring (MRM) mode in negative polarity. Quantification of PIPs was achieved with the MRM pair (Q1/Q3) m/z 413→259. Electrospray (ESI) Voltage was at – 4200 V and TEM (Source Temperature) as 350 °C, DP (Declustering Potential) at -55, EP (Entrance Potential) at -10, CE (Collision Energy) at -31, CXP (Collision cell Exit Potential) at -12. Dwell time of 100 milliseconds was used for experiments with CAD value of -3, GS1 and GS2 at 25, CUR (Curtain gas) at 40. Both Q1 and Q3 masses were scanned at unit mass resolution.

### Total Organic Phosphate measurement

500 µl flow-through obtained from the phosphoinositide binding step of neomycin chromatography was used for the assay for measurements of PI5P. For PI3P measurements, 50 µl was obtained from the last step of lipid extraction and stored separately in phosphate free glass tubes till assay was performed. The sample was heated till drying in a dry heat bath at 90^0^C in phosphate-free glass tubes (Cat# 14-962-26F). The rest of the process was followed as described previously (Jones et al., 2013).

### Software and data analysis

Image analysis was performed by Fiji software (Open source). Mass spec data was acquired on Analyst® 1.6.2 software followed by data processing and visualisation using MultiQuant^TM^ 3.0.1 software and PeakView® Version 2.0., respectively. Chemical structures were drawn with ChemDraw® Version 16.0.1.4. Illustrations were created with BioRender.com. All datasets were statistically analysed using MS-Excel (Office 2016).

## Acknowledgements

This work was supported by the Department of Atomic Energy, Government of India, under Project Identification No. RTI 4006 and a Wellcome-DBT India Alliance Senior Fellowship to PR (IA/S/14/2/501540). We thank the NCBS Imaging and Mass Spectrometry Facility and the *Drosophila* facility for support.

---

**Supporting Figure 1:**
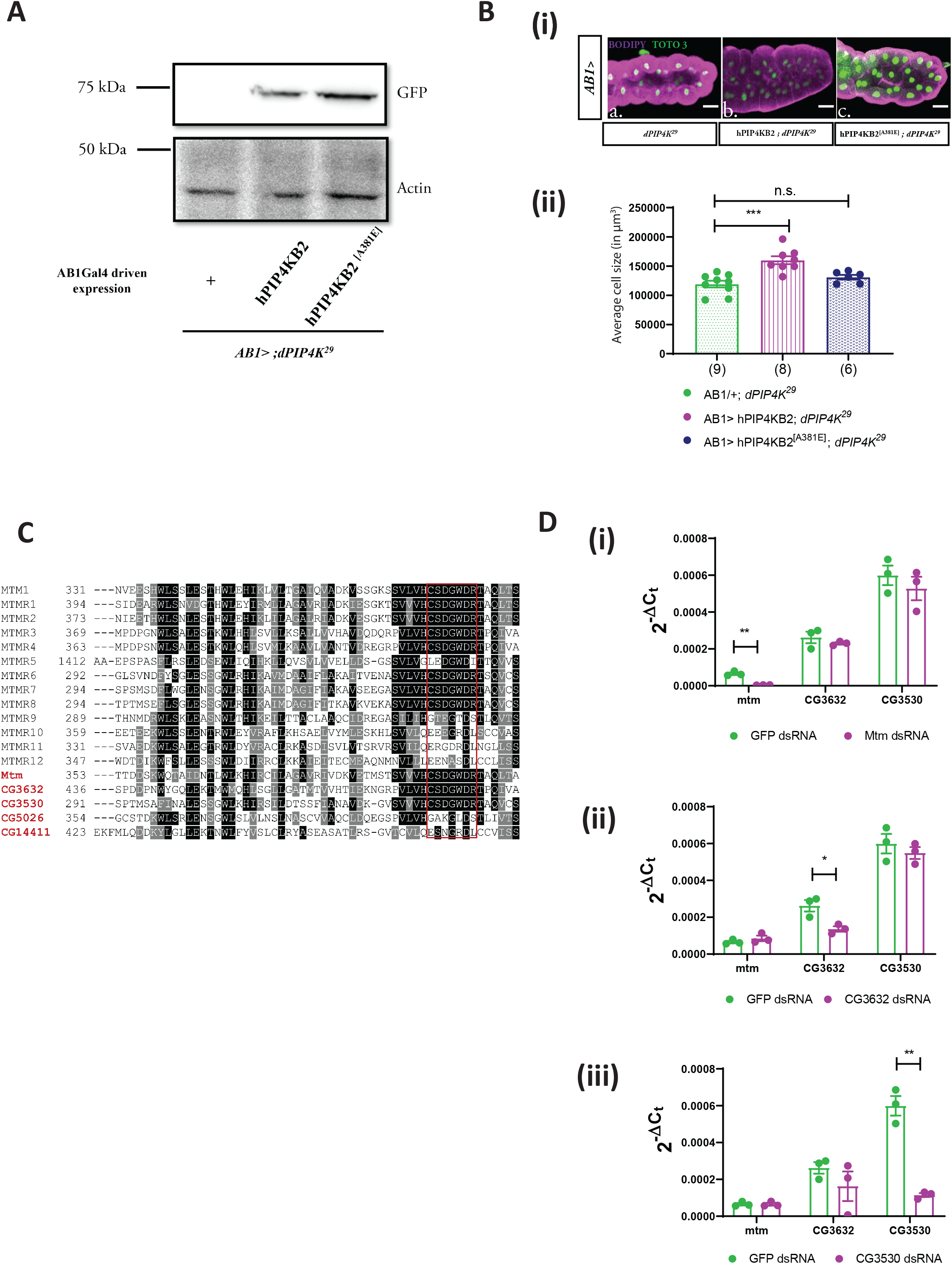
Fab1 and Mtm can are potential PI5P modulating enzymes in *Drosophila*. (A) Protein levels between AB1Gal4/+ ; *dPIP4K^29^* (Ctl), AB1 >hPIP4KB2::GFP ; *dPIP4K^29^* and, AB1 >hPIP4KB2^A381E^::GFP; *dPIP4K^29^* from salivary glands of third instar wandering larvae seen on a Western blot probed by GFP antibody. Both hPIP4KB2::GFP and hPIP4KB2^A381E^::GFP migrates ∼75 kDa. Actin was used as the loading control. (B) (i) Representative confocal images of salivary glands from the genotypes a. *AB1/+ ; dPIP4K^29^*, b. *AB1>hPIP4Kβ; dPIP4K^29^*, c. *AB1>hPIP4Kβ^[A381E]^; dPIP4K^29^*. Cell body is marked majenta by BODIPY conjugated lipid dye, nucleus is marked by TOTO-3 shown in green. Scale bar indicated at 100 μm. (ii) Graph representing average cell size measurement (in μm^3^) as mean ± S.E.M. of salivary glands from wandering third instar larvae of *AB1/+; dPIP4K^29^* (n = 9), *AB1>hPIP4Kβ; dPIP4K^29^* (n = 8), *AB1>hPIP4Kβ^[A381E]^; dPIP4K^29^* (n = 6). Sample size is represented on individual bars. One way ANOVA with post hoc Tukey’s test showed p value = 0.0002 between *AB1/+; dPIP4K^29^* and *AB1>hPIP4Kβ; dPIP4K^29^* and p value = 0.379 between *AB1/+; dPIP4K^29^* and *AB1>hPIP4Kβ^[A381E]^; dPIP4K^29^*. (C) Multiple sequence alignment of the myotubularin phosphatase family proteins in *Drosophila* with human myotubularins share a common signature phosphatase catalytic motif, the C(X_5_)R motif (highlighted in red box) except *CG5026*, *CG14411*. The alignment was generated using clustalO and representation is using Jalview. Conservation is shown in range of white to black, black being most conserved. (D) (i) qPCR measurements for mRNA levels of *Mtm, CG3632* and *CG3530* from either *GFP* ds RNA (green) or *Mtm* ds RNA treated samples (majenta). Student’s unpaired t-test showed p value = 0.009 between *GFP* ds RNA and *Mtm* ds RNA for *Mtm*, while p value = 0.36, p value = 0.46 for genes *CG3632* and *CG3530*, respectively. (ii) qPCR measurements for mRNA levels of *Mtm, CG3632* and *CG3530* from either *GFP* ds RNA (green) or *CG3632* ds RNA treated samples (majenta). Student’s unpaired t-test showed p value = 0.02 between *GFP* ds RNA and *CG3632* ds RNA for *CG3632*, while p value = 0.29, p value = 0.46 for genes *Mtm* and *CG3530*, respectively. (iii) qPCR measurements for mRNA levels of *Mtm, CG3632* and *CG3530* from either *GFP* ds RNA (green) or *CG3530* ds RNA treated samples (majenta). Student’s unpaired t-test showed p value = 0.0008 between *GFP* ds RNA and *CG3530* ds RNA for *CG3530*, while p value = 0.97, p value = 0.31 for genes *Mtm* and *CG3632*, respectively.

**Supporting Figure 2:**
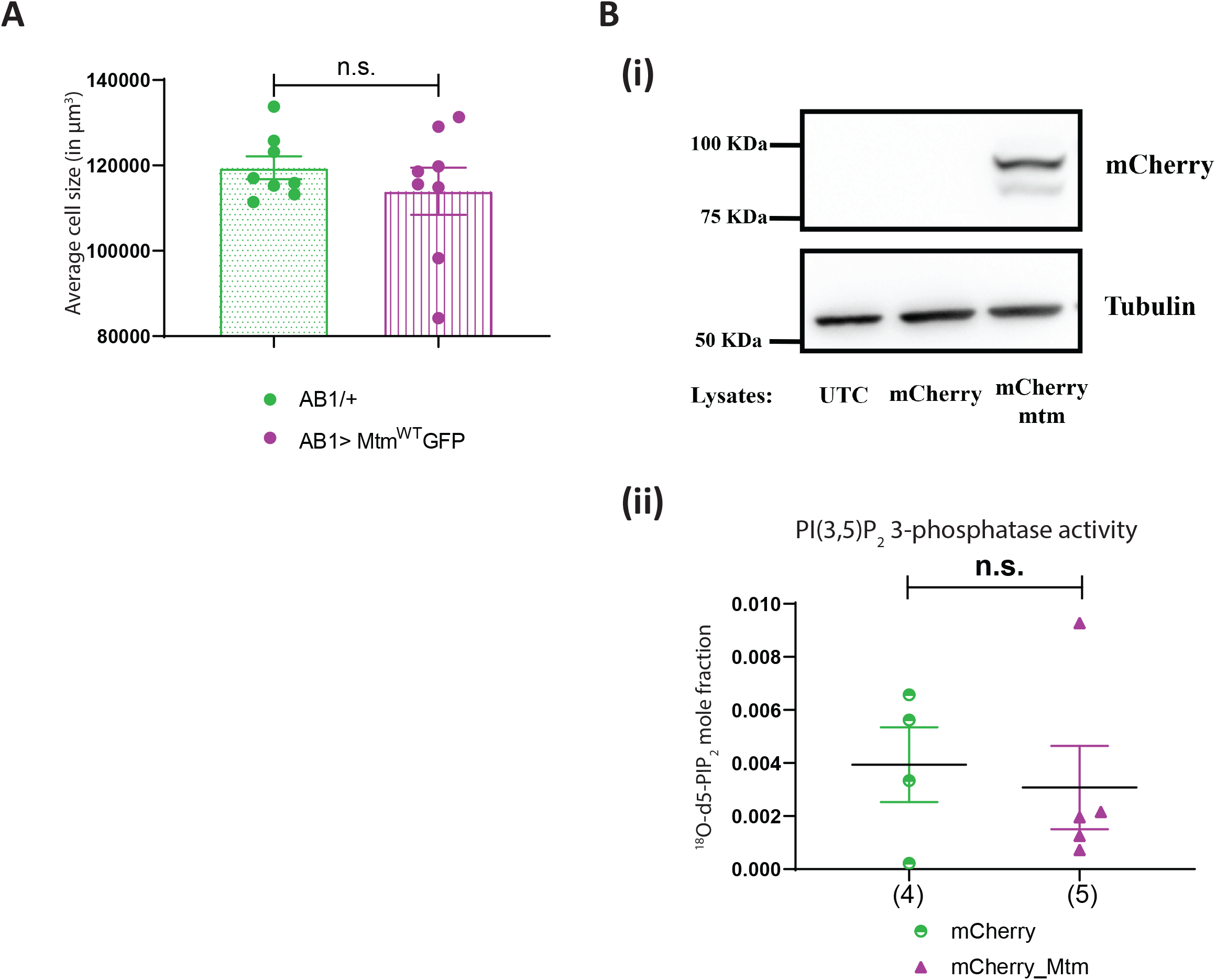
*Drosophila* Mtm rescues the cell size defect of *dPIP4K^29^* independent of PI5P levels. (A) Graph representing average cell size measurement (in percentage) as mean ± S.E.M. of salivary glands from wandering third instar larvae of *AB1/+* (n = 8) and *AB1>*Mtm^WT^GFP (n = 8). Sample size is also represented by points on individual bars. Student’s unpaired t-test with Welch correction showed p value = 0.392. (B) (i) Protein levels between lysates made from *Drosophila* S2R+ cells. Lanes from left: untransfected control (UTC), mCherry vector and mCherry_Mtm observed on a Western blot probed by mCherry antibody. mCherry_Mtm migrates ∼100 kDa. Tubulin was used as the loading control. (ii) *In vitro* phosphatase assay on synthetic PI(3,5)P_2_. Graph representing the formation of ^18^O-PIP_2_ formed from starting PI(3,5)P_2_ as substrate represented as mean ± S.E.M. on addition of either control (mCherry vector transfected lysates) or mCherry_Mtm lysates. Lysate samples n = 3, where each sample was made from five third instar wandering larvae. Student’s unpaired t-test with Welch correction showed p value = 0.696.

**Supporting Figure 3:**
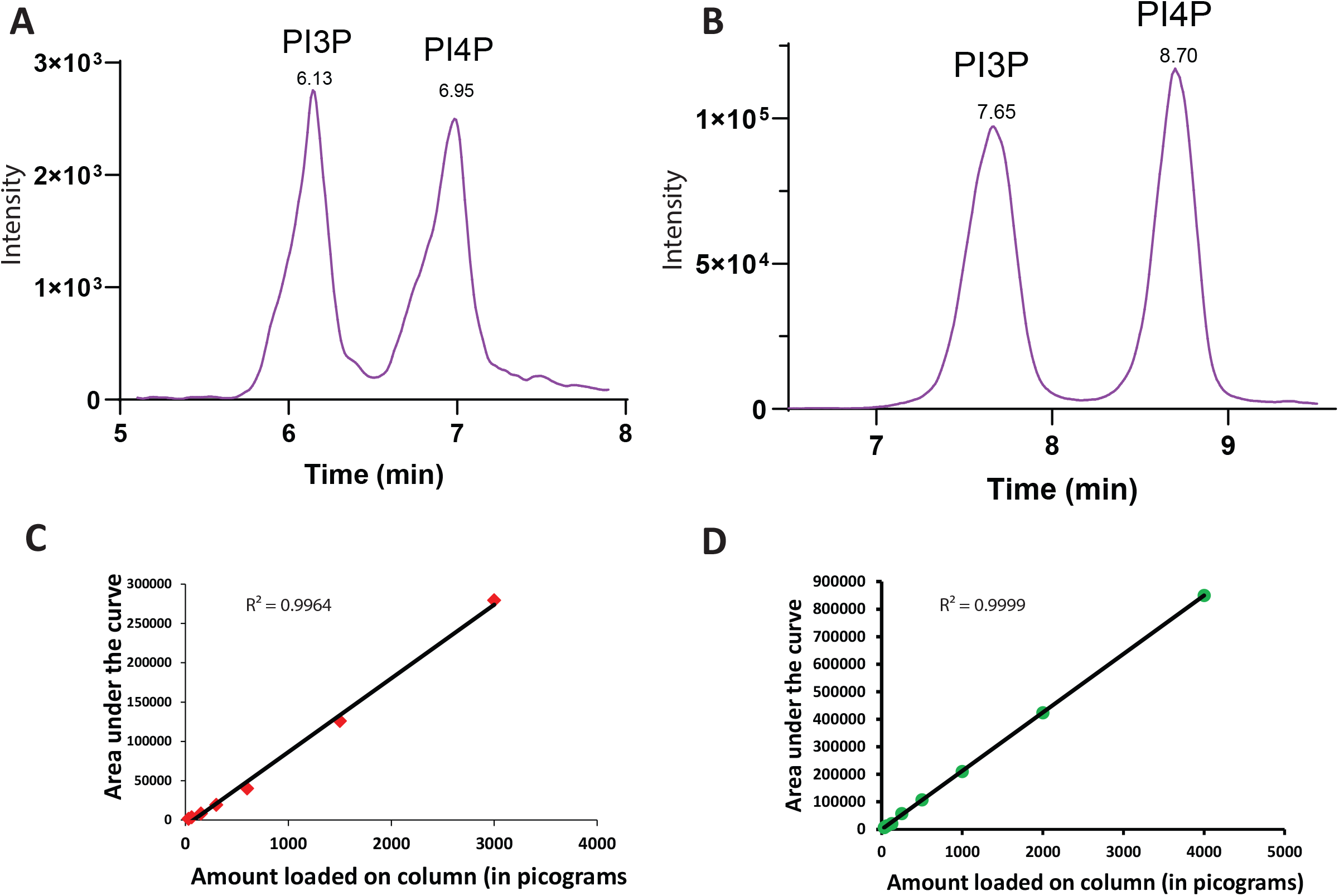
Mtm reduces PI3P levels when over-expressed in *dPIP4K^29^*. (A) XIC obtained from a mixture of synthetic GroPI3P and GroPI4P standard mixture at 300 picograms on column eluting at Rt = 6.13 min and Rt = 6.95 min, respectively. (B) XIC obtained from a biological sample spiked with 200 picograms of synthetic GroPI3P, eluting at Rt = 7.65 min and GroPI4P eluting at Rt = 8.70 min, respectively. The area under the curve (AUC) for GroPI3P changed by 29 times whereas the AUC of GroPI4P changed by 1.5 times, indicating that the first peak obtained in biological samples is indeed GroPI3P. (C) A dose–response curve of synthetic GroPI3P ranging from 30 to 3000 picograms on column. Y-axis depicts intensity of GroPI3P (in cps) and X-axis represents the amount of GroPI3P loaded on column. Equation: y = 93.704x - 7243.1; R² = 0.9964. (D) A dose–response curve of synthetic GroPI4P ranging from 30 to 4000 picograms on column. Y-axis depicts intensity of GroPI4P (in cps) and X-axis represents the amount of GroPI4P loaded on column. Equation: y = 212.49x - 505.82; R² = 0.9999.

**Supporting Figure 4:**
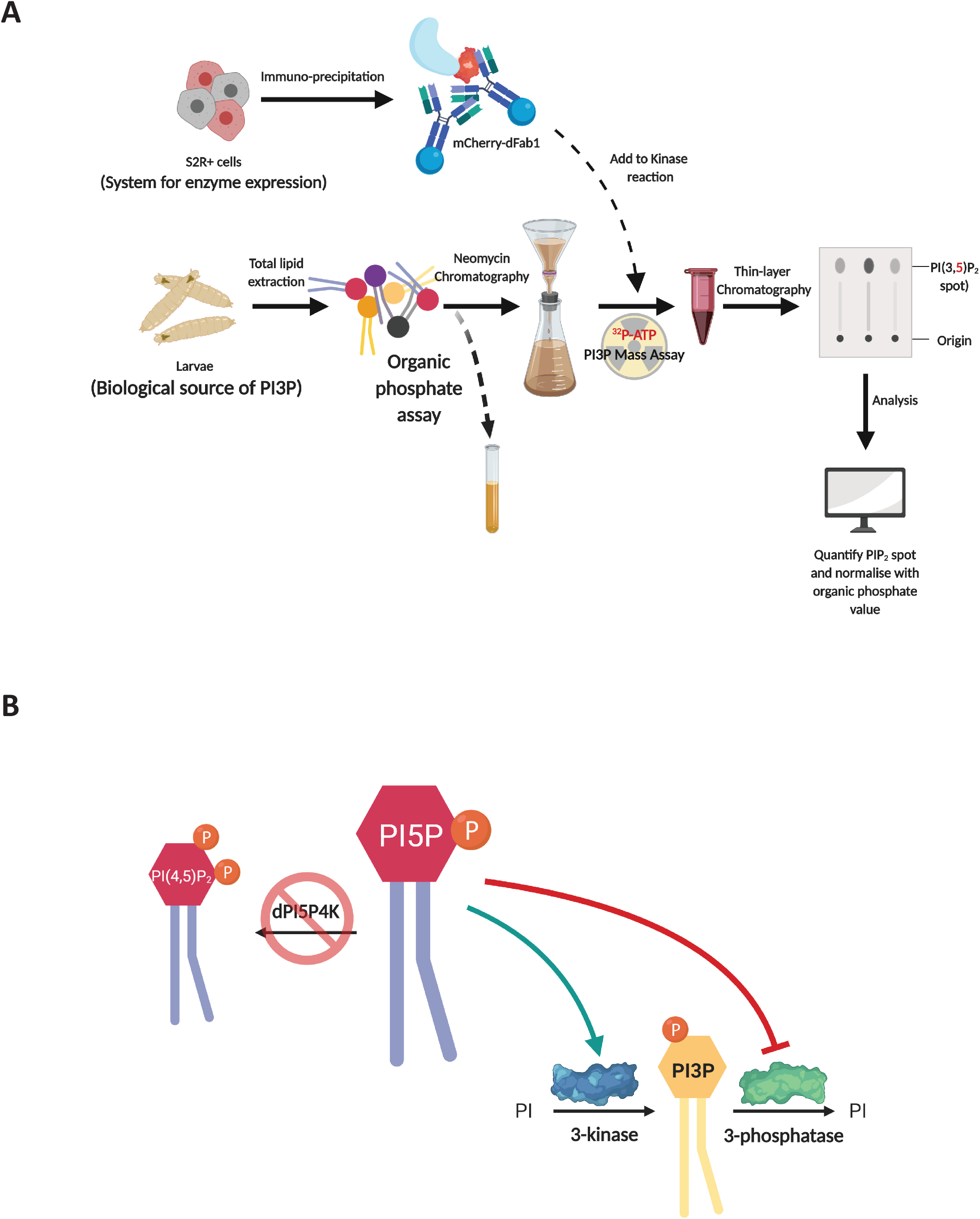
*Drosophila* PIP4K regulates *in vivo* PI3P levels. (A) Schematic illustrating the methodology to assay PI3P by a dFab1 mediated radioactivity-based mass assay. dFab1 is purified from S2R+ cells by immunoprecipitation and used to convert PI3P from total lipid extracts obtained from larvae in presence of ϒ^32^P-ATP to radiolabelled PI(3,5)P_2_ product which is finally analysed using thin layer chromatography (TLC). A portion of the total lipid extract is used for organic phosphate assay to normalise for sample size. (B) Illustration depicting a model where the increased PI3P levels in *dPIP4K^29^* can be explained by either an activation of PI 3-kinase activity (green arrow) or an inhibition of PI3P 3-phosphatase activity (red stubbed arrow).

**Supporting Figure 5:**
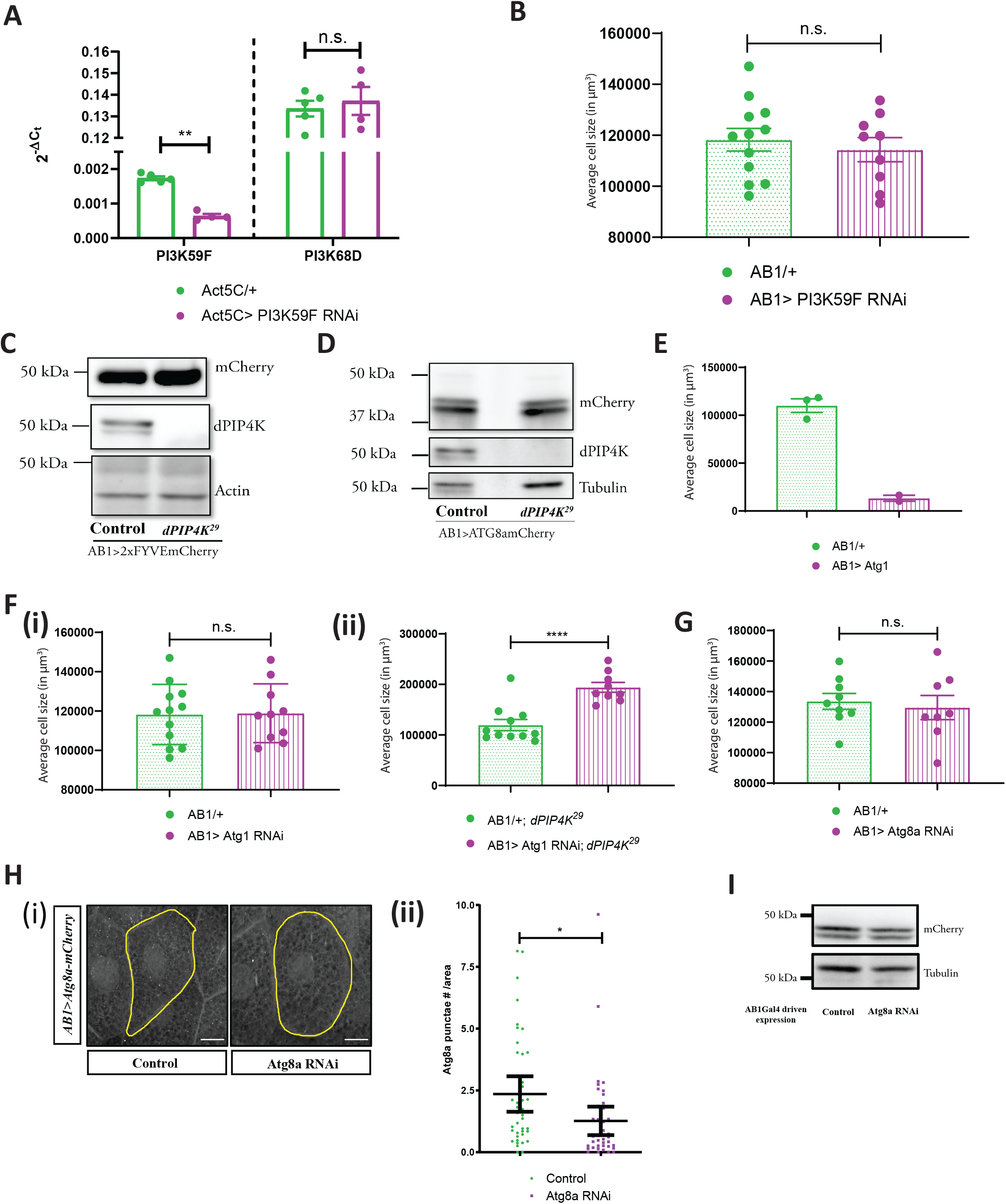
PIP4K in *Drosophila* salivary glands affects bulk autophagy to affect cell size. (A) qPCR measurements for mRNA levels of *PI3K59F* and *PI3K68D* from either Control (Act5C/+, green) or Act5C > *PI3K59F* RNAi (majenta). Multiple t-test with post hoc Holm-Sidak’s test showed p value < 0.0001 between *Act5C/+* and *Act5C> PI3K59F* RNAi for *PI3K59F* and p value = 0.62 between *Act5C/+* and *Act5C> PI3K59F* RNAi for *PI3K68D.* (B) Graph representing average cell size measurement (in μm^3^) as mean ± S.E.M. of salivary glands from wandering third instar larvae of *AB1/+* (n = 12), *AB1>PI3K59F* RNAi (n = 9). Sample size is represented on individual bars. Student’s unpaired t-test with Welch correction showed p value = 0.55. (C) Immunoblot from the salivary glands of wandering third instar larvae probed using mCherry antibody showing expression of 2xFYVE-mCherry in *AB1>2xFYVEmCherry* (control) and *AB1>2xFYVE-mCherry ; dPIP4K^29^*. 2xFYVE-mCherry migrates ∼50 kDa. Actin was used as the loading control. dPIP4K protein was checked in the samples to ascertain the mutant background. (D) Immunoblot from the salivary glands of wandering third instar larvae probed using mCherry antibody showing expression of Atg8a-mCherry in *AB1>ATG8amCherry* (control) and *AB1>ATG8amCherry ; dPIP4K^29^*. Atg8a-mCherry migrates ∼42 kDa. Tubulin was used as the loading control. dPIP4K protein was checked in the samples to ascertain the mutant background. (E) Graph representing average cell size measurement (in μm^3^) as mean ± S.E.M. of salivary glands from wandering third instar larvae of *AB1/+* (n = 3), *AB1> Atg1 (*n = 2). Sample size is represented on individual bars. Statistical test not performed. (F) (i) Graph representing average cell size measurement (in μm^3^) as mean ± S.E.M. of salivary glands from wandering third instar larvae of *AB1/+* (n = 12), *AB1> Atg1* RNAi (n = 10). Sample size is represented on individual bars. Student’s unpaired t-test with Welch correction showed p value = 0.92. (ii) Graph representing average cell size measurement (in μm^3^) as mean ± S.E.M. of salivary glands from wandering third instar larvae of *AB1/+ ; dPIP4K^29^* (n = 11), *AB1>Atg1* RNAi*; dPIP4K^29^* (n = 9). Sample size is represented on individual bars. Student’s unpaired t-test with Welch correction showed p value <0.0001. (G) Graph representing average cell size measurement (in μm^3^) as mean ± S.E.M. of salivary glands from wandering third instar larvae of *AB1/+* (n = 9), *AB1>Atg8a* RNAi (n = 8). Sample size is represented on individual bars. Student’s unpaired t-test with Welch correction showed p value = 0.67. (H) (i) Representative confocal z-projections depicting autophagosomal levels using Atg8a-mCherry in the salivary glands from the genotypes a. *AB1>ATG8a-mCherry,* b. *AB1>ATG8a-mCherry ; ATG8aRNAi* . Scale bar indicated at 20 μm. (ii) Graph representing Atg8a punctae measurement in the salivary glands from wandering third instar larvae of *AB1>ATG8a-mCherry* (N =8, n =40) and *AB1>ATG8a-mCherry ; ATG8aRNAi* (N =8, n =40). Student’s unpaired t-test with Welch correction showed p value = 0.0197. (I) Immunoblot from the salivary glands of wandering third instar larvae probed using mCherry antibody showing the expression of Atg8a-mCherry in *AB1>ATG8amCherry* (control) and *AB1>ATG8a-mCherry ; ATG8aRNAi*. Atg8a-mCherry migrates ∼42 kDa. Actin was used as the loading control.

**Supporting Figure 6:**
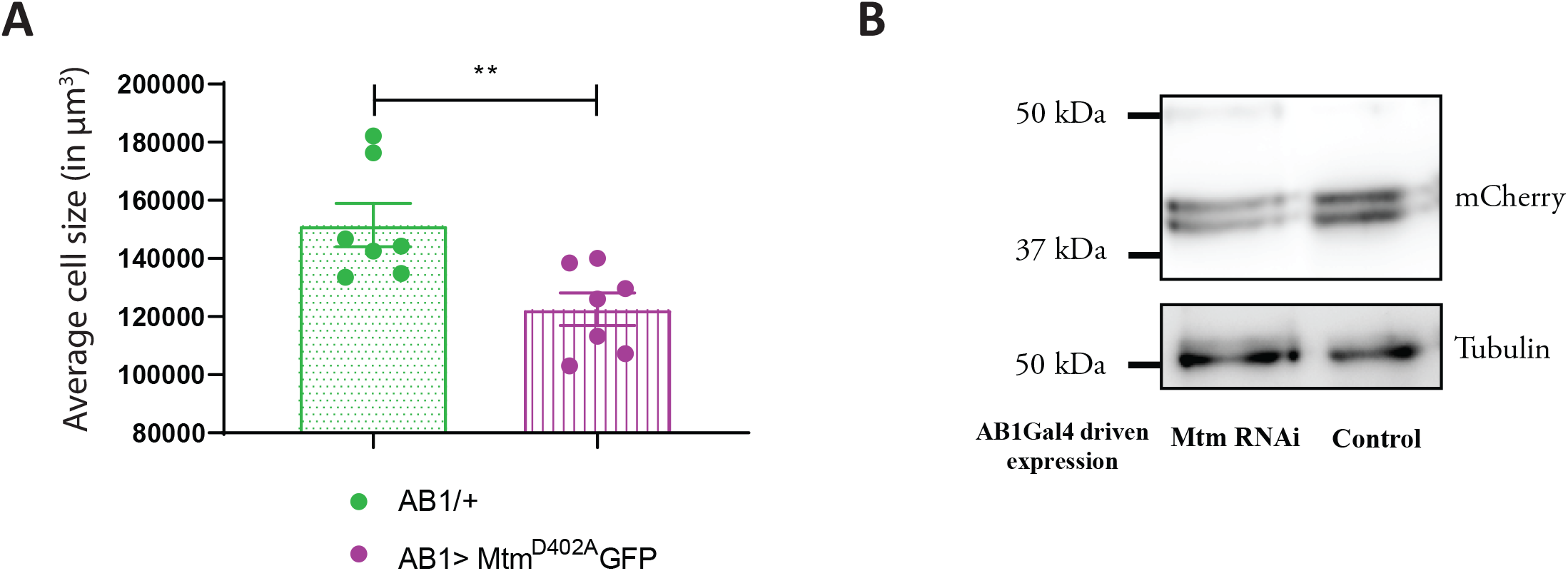
PI3P regulates cell size in salivary glands. (A) Graph representing average cell size measurement (in μm^3^) as mean ± S.E.M. of salivary glands from wandering third instar larvae of *AB1Gal4/+* (n = 7), *AB1> Mtm* RNAi (n = 7). Sample size is represented on individual bars. Student’s unpaired t-test with Welch correction showed p value = 0.009. (B) Immunoblot from the salivary glands of wandering third instar larvae probed using mCherry antibody showing the expression of Atg8a-mCherry in *AB1>ATG8a-mCherry* (control) and *AB1>ATG8a-mCherry; Mtm RNAi*. Atg8a-mCherry migrates ∼42 kDa. Tubulin was used as the loading control.

